# Differential effects of carbon nanotube and graphene on the tomato rhizosphere microbiome

**DOI:** 10.1101/2022.11.10.516042

**Authors:** Yaqi You, Patricia Kerner, Sudha Shanmugam, Mariya V. Khodakovskaya

**Affiliations:** Department of Environmental Resources Engineering, SUNY College of Environmental Science and Forestry, Syracuse, NY, USA; Department of Biological Sciences, Idaho State University, Pocatello, ID, USA; Department of Biology, University of Arkansas at Little Rock, Little Rock, AR, USA

**Keywords:** nanomaterials, carbon nanotube, graphene, tomato, microbial community, soil microbiome, rhizosphere, metagenomics

## Abstract

Application of carbonaceous nanomaterials (CNMs) to the soil-plant system can affect plant physiology, with positive results ranging from enhanced seed germination and root system development to improved stress tolerance. The underlying mechanisms are not fully understood. Plant rhizosphere microbiomes at the soil-root interface are strongly influenced by the host plant and play a key role in the plant host’s development and health. Yet few studies have characterized changes in plant rhizosphere microbiomes following applications of CNMs to the soil-plant system. Here we investigated the effects of multi-walled carbon nanotube (CNT) and graphene on microbial communities in the ectorhizosphere of tomato plants versus surrounding bulk soil. Pot experiments were conducted where tomato plants were exposed to CNT or graphene at 200 mg/kg soil for four weeks. Ectorhizosphere and bulk soils were then collected and analyzed for physicochemical properties and microbiome structure and function. While graphene had a limited impact on the tomato rhizosphere microbiome, CNT significantly increased microbial alpha diversity, induced greater divergence of beta diversity, enhanced microbial interactions, and potentially impacted community functions such as aromatic compound degradation, antioxidant synthesis, and redox cofactor biosynthesis. Furthermore, CNT induced stronger and/or unique microbiome alterations in the tomato rhizosphere compared to bulk soil. Our findings reveal the differential modulating effects of two widely-used CNMs on plant rhizosphere microbiomes and highlight an imminent need to understand complex plant root-microbe interplays in the CNM-impacted rhizosphere. These results have implication for realizing the full potential of phytoapplication of CNMs toward improved and sustainable plant production.

## Introduction

The current food and agricultural system faces significant sustainable challenges due to rapid population growth, increasing global food demand and inefficient utilization of resources, and left a huge ecological footprint on the Earth. Nanotechnology is promising to offer innovative solutions to many of these challenges due to its improved capabilities to sense and monitor physical, chemical or biological processes, slow-release fertilizers or nutrients with time, control microbes associated with animal/plant hosts, and mediate biomolecule delivery for genetic engineering of plants.^1-4^ Of increasing interest is phytoapplication of carbonaceous nanomaterials (CNMs).^5-8^ They have been shown to enhance seed germination, stimulate plant reproductive system, increase shoot and root biomass production, activate photosynthesis, boost phytomedicinal contents, and improve plant tolerance and crop yield under salinity and drought stresses.^9-18^

While positive plant phenotypic changes upon CNM application have long been recognized, the underlying mechanisms are not fully understood. Khodakovskaya et al. (2011) reported the uptake of multi-walled carbon nanotubes (CNT) by tomato plants and translocation of CNT to roots, leaves, and fruits, which resulted in the activation of the aquaporin (*LeAqp2*) gene in roots and many stress-signaling genes (e.g., heat shock protein 90 or HSP90 gene) in various tissues, concequently enhancing germination and growth of tomato seedlings.^19^ Cao et al. (2020) found that in tomato, multi-walled CNT induced higher nitrate reductase activity and thus endogenous nitric oxide (NO) production, which in turn stimulated lateral root development, likely through NO signaling.^20^ Stimulated seed germination and plant growth was observed after the application of not only CNT with different lengths and shapes but also a wide range of other CNMs including graphene and nanohorns.^14,21^ Transcriptomics analysis revealed that graphene induced up-regulation of transcriptional factor, plant hormone signal transduction, nitrogen and potassium metabolism, and secondary metabolism in maize roots.^22^ More recently, Rezaei Cherati et al. (2021) found that CNT and graphene enhanced tolerance to salinity and drought in tomato, rice, and sorghum by reconciling the expression of affected stress-responsive genes, including those encoding transcription/translation factors, dehydrins, heat shock proteins and aquaporins, and genes involved in calcium transport, mitogen-activated protein kinase (MAPK) signaling cascade, abscisic acid metabolism and signaling.^18^ The same study further found that CNT restored more genes than graphene in tomato seedlings under salinity stress, while graphene restored more genes than CNT in rice under drought stress. At the metabolome level, CNT applied to soil as plant growth regulators affected tomato pathways including carbon metabolism, biosynthesis of amino acids and secondary metabolites, amino sugar and nucleotide sugar metabolism, among others.^23^ Moreover, foliar-applied carbon dots were shown to promote maize drought tolerance by increasing the levels of root exudates (e.g., succinic acid, pyruvic acid, and betaine).^8^

Besides modulating plant biological pathways, CNMs can influence the soil microbiome through changing microbial abundance, diversity, community composition, and function. To date, both positive and negative effects of CNMs have been reported, depending on the type of nanomaterial, the application rate, and the exposure duration.^10,24-26^ Soil microbes, especially those in the rhizosphere, play a key role in the development and health of plant systems.^27-31^ Together they govern nutrient cycling, mediate plant-beneficial functions such as disease suppression, and form holobionts with the plant hosts.^32^ Therefore, many of the CNM-introduced plant phenotypic changes might be associated with plant-associated microbiomes. Yet few studies have elucidated shifts in the rhizosphere microbiome following phytoapplication of CNMs, nor has research addressed whether the shifts are comparable to those occurring in the bulk soil.

This study aimed to bridge this gap by investigating the effects of two widely-used CNMs, CNT and graphene, on microbial communities in the rhizosphere and surrounding bulk soil of tomato plants, one of the most consumed vegetable crops around the world. Specifically, we focused on the ectorhizosphere, the outermost zone that extends from the rhizoplane next to the root epidermal cells and mucilage into the bulk soil, because plant-microbe interactions in this zone strongly influence the host itself.^30,31^ We hypothesized that the two CNMs would (1) modulate the structure and function of tomato-associated microbiomes differentially, and (2) induce unique microbiome alterations in the tomato rhizosphere that did not occur in the bulk soil. Addressing how various nanomaterials modulate the plant rhizosphere microbiome not only adds to the understanding of the mechanism underlying nanomaterial-introduced plant phenotypic changes but may also shed light on harnessing targeted manipulation of the rhizosphere microbiome using various nanomaterials.

## Materials and Methods

### Preparation and characterization of CNMs

Surface functionalized (-COOH) multi-walled CNT and graphene nanoplatelet were purchased from Cheap Tubes (Brattleboro, VT, USA). Multi-walled CNT had an OD of 13−18 nm and a length of 1−12 μm. Graphene had a lateral dimension of 1-2 μm with <3 layers. CNMs were characterized using transmission electron microscopy (TEM) and Raman spectroscopy as before.^14,33^ CNMs were suspended in sterile MilliQ water and the suspensions were autoclaved three times at 121°C, 15 lb/in^2^ for 20 min to eliminate potential endotoxin contamination as described by Lahiani et al. (2016).^14^

### Plant cultivation and CNM exposure experiments

A model cultivar of tomato (*Solanum lycopersicum* L.), Micro-Tom, was used in this study. Tomato seeds obtained from Reimer Seeds Co. Inc. (Saint Leonard, MD, USA) were sterilized as before.^18^ Sterile seeds were placed in small germination pots and incubated in a growth chamber for 21 days with the following conditions: 24 °C, 12 hours of light with a light intensity of 105 μmol/s m^2^. At the end of the germination period, 3-week-old tomato seedlings were transferred to experimental pots containing ∼400 g of Sungro Horticulture growing soil mix for exposure experiments.

CNM exposure experiments were conducted in a greenhouse for 3 weeks under the following conditions: 8-h light (26 C°), 16-h dark (22 C°). In the CNT experiment, six control plants received 100 mL of deionized water weekly while another 6 plants received 100 mL of CNT solution (200 μg/mL) weekly for 3 weeks. Similarly, in the graphene experiment, five control plants received 100 mL of deionized water weekly while another 6 plants received 100 mL of graphene solution (200 μg/mL) weekly for 3 weeks. The final amount of either CNM in each pot was approximately 150 mg/kg of soil mix. This concentration was selected because it was previously found as the most effective nanofertilization concentration for tomato plants.^13^ The selected dose was also comparable to those used in prior studies of CNM effects on the soil-plant system (Chen et al. 2022),^24-26,34-36^ although higher than estimated soil concentrations of CNMs.^37,38^

### Soil sample collection and processing

Bulk and rhizosphere soils were collected separately from each of the 23 pots after the exposure experiment (10 week-old plants), using approaches similar to others.^39,40^ First, the tomato plant was carefully removed from the pot using aseptic technique. Loosely bound soil was removed from the roots by shaking and using a sterilized spatula. The roots with tightly bound soil (2-5 mm thick) were carefully transferred into a sterile 50 mL centrifuge tube, immediately frozen in liquid nitrogen, and stored at -80 °C until further processing. For DNA extraction, 15 mL autoclaved, cold 1X phosphate-buffered saline (pH 7.4) and 15 μL Triton X-100 (0.22 μm filtered) were added to each 50 mL centrifuge tube containing roots with tightly bound soil. After gently vortexing, clean roots with minimal soil attached to the surface were carefully removed from the 50 mL centrifuge using sterilized tweezers. The remaining suspension was centrifuged at 4 °C, 6000x g for 10 min. The resulting supernatant was discarded, and the soil pellet was used in DNA extraction as detailed below.

For each processed experiment pot, soil in close proximity to the plant’s root system was removed using a sterilized spatula and any small root in the remaining soil was removed with sterilized tweezers. The remaining bulk soil was then mixed using the sterilized spatula, transferred into a sterile 50 mL centrifuge tube, immediately frozen in liquid nitrogen, and stored at -80 °C until DNA extraction. A second set of mixed bulk soil was used in community-level functional analyses (EcoPlate assay and basal soil respiration test) as detailed below. Additional mixed bulk soil was archived for soil physicochemical property analyses (details in the SM).

### Assessing substrate use pattern and basal respiration of the soil microbial community

Due to the limited amount of rhizosphere soils, only bulk soils were measured for community-level microbial functionality. Six bulk soils from the CNT experiment and eight bulk soils from the graphene experiment were assessed for substrate use pattern with EcoPlates (Biolog, Hayward, CA).^41^ Each EcoPlate device is a 96-well plate containing a range of substrate sources prepared in triplicate. Bulk soil was thoroughly mixed, suspended in sterile Milli-Q water (1:10 w/v), dispersed for 10 min in a FS20 ultrasonic bath (Fisher Scientific, Hampton, NH), and shaken in an incubator at room temperature, 250 rpm for 20 min. To prevent interference from soil nutrients while maintaining sufficient inoculant microbes, the soil suspension was diluted by 1:5000, and 150 μL of the diluted suspension was added to each well in an EcoPlate. The plate was incubated at 25 °C in the dark under a humid condition, and color development was monitored by measuring optical density at 590 nm (OD_590_) at an interval of approximately 24 hours for ∼170 hours using a microplate reader (Synergy HTX, Biotek Instruments, Winooski, VT). Microbial activity in each microplate, expressed as average well-color development (AWCD), was determined as follows: AWCD = Σ(OD_590,i_/31) where OD_590,i_ is the optical density value from each well and 31 represents the 31 sole substrate sources in each plate. The final reading was compared across technical replicates (n = 3) and pot replicates (n = 3∼4). Soil basal respiration was measured to assess soil microbial activity, using an approach similar to others.^34^ More details about EcoPlate assay and soil respiration measurement are presented in the SM.

### DNA extraction and 16S sequencing

Total DNA was extracted from bulk (n = 23) and rhizosphere (n = 23) soils using the PowerSoil DNA Kit or the PowerSoil Pro Kit (QIAGEN) following the manufacturer’s protocols with minor modifications. Briefly, when the PowerSoil DNA Kit was used, the inhibitor removal step was conducted at 4 °C for 10-20 min. A PowerLyzer homogenizer was used to efficiently grind samples with reduced heat generation. Extracted DNA was further purified using the ethanol precipitation protocol.^42^ The quantity and quality of the resulting DNA extractants were evaluated using a NanoDrop 1000 spectrophotometer (Thermo Fisher Scientific) as well as by gel electrophoresis.^43^ All the DNA samples were stored at -20 °C before further processing.

Library preparation was conducted similarly to the Earth Microbiome project and as before.^43-45^ The V4 region of prokaryotic 16S rRNA gene (∼390 bp) was amplified with forward-barcoded 515F primer (5’-3’: GTGYCAGCMGCCGCGGTAA) and 806R primer (5’-3’: GGACTACNVGGGTWTCTAAT).^46-48^ A typical PCR reaction consisted of 20 ng of template DNA in 1 μL, 0.2 μM of each primer, 12 μL of the Invitrogen Platinum Hot Start PCR Master Mix (Thermo Fisher Scientific), and nuclease free water in a total volume of 25 μL. Libraries were cleaned up with the MagBio HighPrep PCR Clean-up System (Illumina), quantified on a CFX96 Touch Real-Time PCR System (Bio-Rad) with the NEBNext Library Quant Kit for Illumina (New England BioLabs), and checked for quality on an 5200 Fragment Analyzer (Agilent Technologies) utilizing the High Sensitivity NGS Fragment Kit (Agilent Technologies). After quantification on a Qubit 4.0 fluorometer (Thermo Fisher Scientific) with the QuantiFluor dsDNA HS System (Promega), the libraries were pooled in equimolar concentrations and sequenced on an Illumina MiSeq platform with the MiSeq Reagent Kit v2 (500-cycles; Illumina). PhiX control v3 was used to ensure a 3-5% spike.

### Bioinformatics and statistical analysis

Paired-end reads (2×250 bp) were subjected to quality control with FastQC (Andrews 2017) and then processed using the QIIME2 pipeline (version 2021.2) (Bolyen et al. 2019).^49,50^ Briefly, raw reads were trimmed at both ends when mean Phred values dropped below 30 and denoised using DADA2 to revolve amplicon sequence variants (ASVs).^51^ This approach allows one to resolve one-nucleotide differences.^52^ Rare ASVs (<10 occurrence among all the samples) were filtered out to avoid sequencing errors and artifacts. Prokaryotic taxonomy was assigned using a naive Bayes machine-learning classifier that was trained for the 515F-806R V4 region against the SILVA database (release 138).^53,54^ Reads associated with Eukarya, mitochondria and chloroplast, as well as unassigned reads including those without a defined Phylum, were filtered out. For comparison purposes, ASVs present in at least 2 samples from the CNT or graphene experiments (68% and 86% reads, respectively) were retained. This resulted in 146426-336538 reads (median 252095) per sample among all the soils (n = 46).

Community alpha diversity was calculated at the ASV level after rarefaction. Community beta diversity was analyzed using log transformed ASV abundance and visualized with principal coordinate analysis (PCoA) of Bray-Curtis and weighted UniFrac distance.^55^ Microbial network analysis was performed using Bray-Curtis distance of agglomerated ASV abundance to the phylum level (maximum distance set to 0.3). Linear discriminant analysis (LDA) of effect size (LEfSe) was applied to ASV relative abundance to identify differential taxa that were most likely to explain differences between samples. For a taxonomy to be considered as discriminative, the threshold on the absolute value of the logarithmic LDA was set to 2 and the alpha values for the ANOVA and Wilcoxon tests were set to 0.05.^56^

To profile putative microbial functions, we used PICRUSt2 (Phylogenetic Investigation of Communities by Reconstruction of Unobserved States) which expands PICRUSt original database of gene families and reference genomes (>20 fold increase in reference genomes from the IMG (Integrated Microbial Genomes) database), and is thus more accurate with reduced bias as rare environmental-specific functions can be more readily predicted.^57^ Gene family abundance was inferred based on Kyoto Encyclopedia of Genes and Genomes (KEGG) orthologs (KOs) and Enzyme Commission numbers (EC numbers). Pathway abundance was inferred based on the MetaCyc database through structured mappings of EC gene families. The nearest-sequenced taxon index (NSTI) was calculated for each input ASV and any ASV with NSTI > 2 was excluded from the output. Pathway enrichment was identified based on the PICRUSt2 results using DESeq2 and MicrobiomeAnalyst.^58,59^

Metagenomic data analysis and visualization were implemented using R (version 4.0.3) with the package ‘tidyverse’, ‘ggplot2’, ‘igraph’, ‘phyloseq’, ‘phyloseqGraphTest’, ‘DESeq2’, ‘vegan’, ‘ggnetwork’, among others.^60-64^ Significant differences in alpha and beta diversity were determined using the nonparametric Kruskal-Wallis test and permutational multivariate analysis of variance (PERMANOVA), respectively.^65,66^ To test the influence of treatment or soil zone on the microbial network, graph-based permutation test was conducted using the minimum spanning tree (n = 9999) and the nearest neighbor (NN) tree (knn = 1).^67^ If two nodes (phyla) were of the same treatment condition or soil zone, the edge connecting them was “pure”; otherwise, the edge was ‘‘mixed.’’

### Availability of Sequence Data

All Illumina reads are deposited at the NCBI Sequence Read Archive (SRA), under the BioProject accession number PRJNA900043 (https://www.ncbi.nlm.nih.gov/bioproject/PRJNA900043).

## Results and Discussion

### CNMs changed bulk soil properties

The inclusion of corresponding controls allowed us to identify changes due to CNM treatment. CNT increased bulk soil electrical conductivity (EC) by 74%, NO_3_-N by 21%, SO_4_-S by 250%, Zn by 26%, and Cn by 27%, while decreasing CaCO_3_ by 59%, Fe by 50%, and Mn by 32%. Graphene increased bulk soil organic matter (OM) by 30%, total C by 27%, C/N ratio by 20%, and CaCO_3_ by 52%, while decreasing P by 37% (Table S1). In addition, CNT resulted in reduced cation exchange capacity (CEC) and NH_4_-N, whereas graphene enhanced these soil properties. Interestingly, neither nanomaterial influenced soil pH, which remained slightly acid, nor did either CNM affect total N in the soil. These soil physicochemical changes may have resulted directly from the addition of CNMs. CNT was reported to contain higher metal contents than graphene.^68^ Both CNMs have high metal adsorption capacity and can immobilize metals in soil.^69^ Alternatively, the soil physicochemical changes may be a result of alterations in the soil-plant system (e.g., update/release of nutrients by plant, soil enzyme activity). For example, soil amendment with 200 mg/kg CNT was reported to increase corn K update and the activity of urease and dehydrogenase in soil, but decreased phosphatase activity.^36^ Regardless of the exact mechanism, soil physicochemical property changes could in turn influence microorganisms living in the soil environment.

### CNT significantly increased microbial diversity in the tomato rhizosphere

CNT increased microbial richness in bulk soil and the tomato rhizosphere (Figure 1). The average ASVs were 3718 and 3376 for bulk and rhizosphere soils from the control pots, respectively, and 4981 and 4493 ASVs for bulk and rhizosphere soils from the treatment pots, respectively. Similarly, graphene increased microbial richness in bulk soil and the tomato rhizosphere. The average ASVs were 4341 and 3995 for bulk and rhizosphere soils from the control pots, respectively, and 4600 and 4378 for bulk and rhizosphere soils from the treatment pots, respectively. Considering both soil zones, soils from the CNT experiment had a smaller core soil microbiome (27% of all ASVs) than soils from the graphene experiment (51% of all ASVs) (Figure 1).

**Figure 1.**
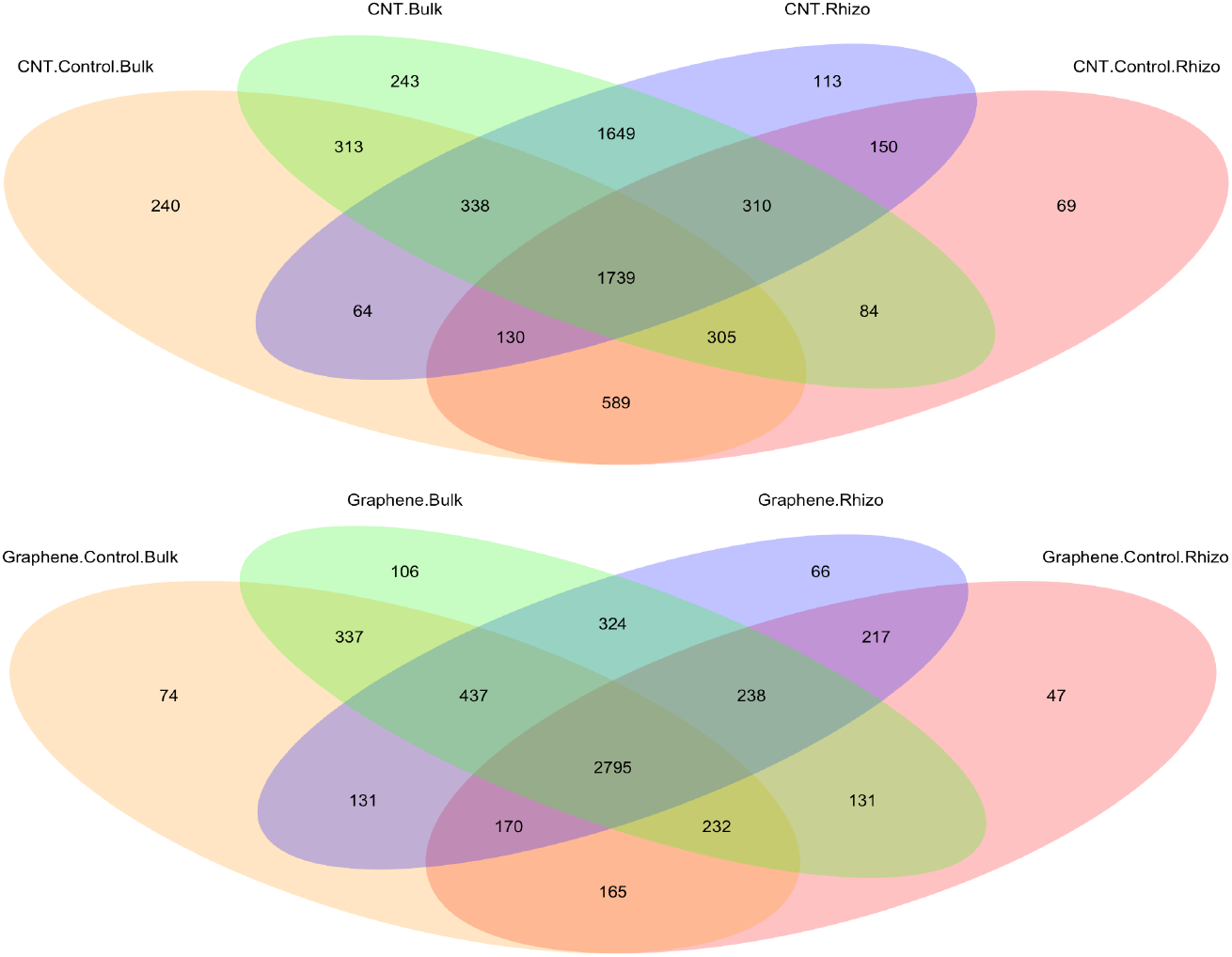
Venn diagram showing ASVs identified in the soils receiving different treatments.

Changes in microbial alpha diversity were further manifested in Shannon index (Figure 2). CNT increased Shannon diversity in both bulk soil and the tomato rhizosphere, with the increase in the rhizosphere being significant (from 5.92 to 6.33, *p* = 0.037 in Kruskal-Wallis test). Consequently, despite the control pots having higher Shannon diversity in bulk soil than the rhizosphere (*p* = 0.006, Kruskal-Wallis test), the CNT treatment pots showed no significant difference in Shannon diversity between the two soil zones (*p* = 0.200, Kruskal-Wallis test). In the graphene experiment, bulk and rhizosphere soils had no significant difference in microbial Shannon diversity, and graphene did not affect microbial Shannon diversity in either soil zone.

**Figure 2.**
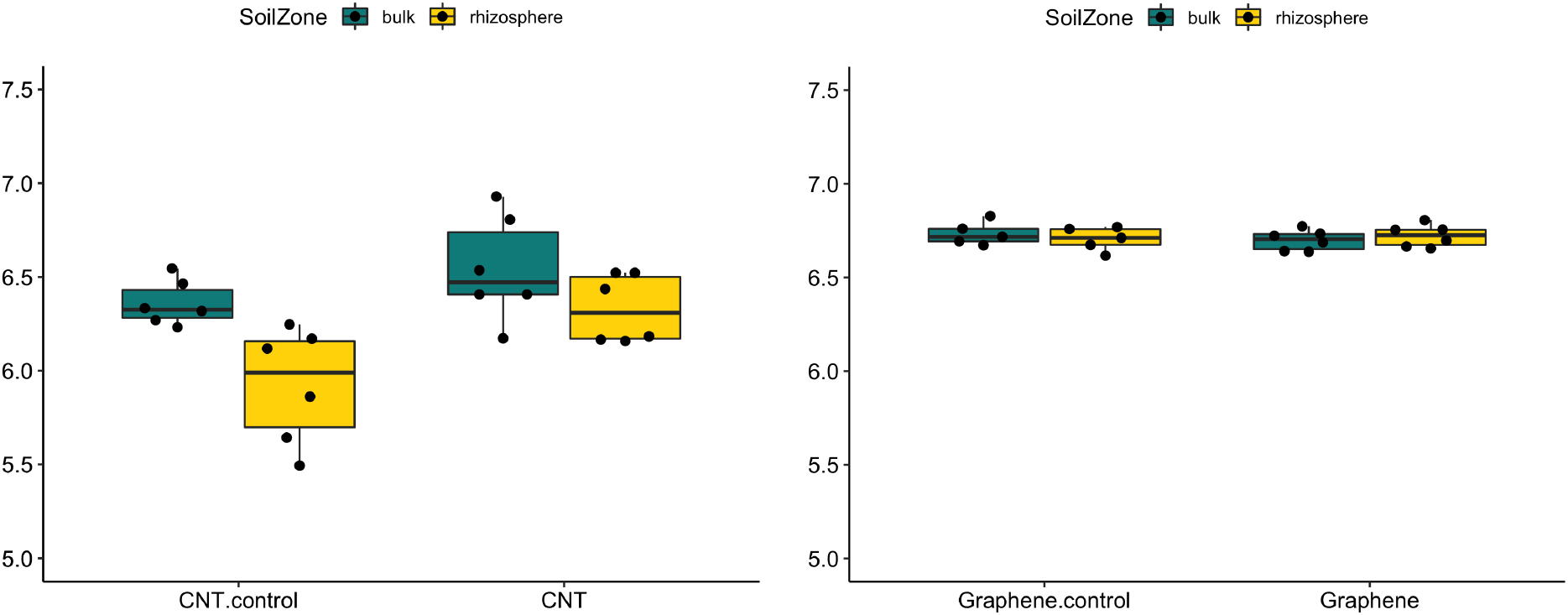
Effects of CNT (left) and graphene (right) on microbial Shannon diversity in bulk soil and the tomato rhizosphere.

Diversity is one of the most commonly monitored microbial community indices when studying the influence of CNMs on the soil microbiome, as drastic reduction in microbial diversity could suggest degraded soil health. Prior research has unveiled mixed effects of CNMs, depending on the type of nanomaterial, the application rate, and the exposure duration. One earlier study found that multi-walled CNT had no significant effect on bacterial diversity in tomato-growing soil.^13^ In another study, both single-walled and multi-walled CNT stimulated bacterial alpha diversity in bulk soil.^24^ Fewer studies have investigated the influence of CNMs on microbial diversity in the rhizosphere. A recent study of *Solanum nigrum* found no significant effect of multi-walled CNT on the alpha diversity of rhizosphere bacterial or fungal communities.^26^ Here, our results suggest that CNT but not graphene influenced soil microbiomes substantially, with a greater impact on the tomato rhizosphere than bulk soil. While these observations could be partially attributed to CNT-introduced soil property changes, a stronger shift in the rhizosphere suggests that specific conditions in that microenvironment, such as CNM-induced root exudates, may have contributed to the assembly of a differential rhizosphere microbiome compared. Indeed, metabolomics analysis revealed that soil amendment of CNT could induce up- or down-regulation of a variety of pathways in tomato roots, including carbon metabolism, biosynthesis of amino acids and secondary metabolites, amino sugar and nucleotide sugar metabolism, among others.^23^ Similarly, foliar application of carbon dots to maize seedlings increased the levels of succinic acid, pyruvic acid, and betaine in root exudates.^8^ It is very likely that in this study, CNMs triggered changes in the tomato root metabolome. Future multi-omics analyses integrating plant metabolomics and microbial metagenomics are needed to elucidate mechanisms underlying CNM-induced interplay between root exudates and the rhizosphere microbiome.

### CNT induced greater divergence of the tomato rhizosphere microbiome

The plant rhizosphere microbiome, shaped by plant-microbe coevolution, is known to differ from the bulk soil microbiome (Trivedi et al. 2020). Here we observed distinct microbial communities in bulk soil and the tomato rhizosphere for all the experiment pots (*p* < 0.005 in PERMANOVA test) (Figure 3). CNT significantly altered microbial beta diversity in both soil zones (*p* < 0.001 in PERMANOVA test) (Figure 3), such that in the PCoA plot, soils separated from each other by treatment along the primary coordinate (explaining 26% of the total variance) and by soil zone along the secondary coordinate (explaining 14% of the total variance). In contrast, graphene did not affect microbial beta diversity in either soil zone (*p* > 0.1 in PERMANOVA test) (Figure 3). In the PCoA plot, soils from the graphene experiment are separated primarily by soil zone along the first coordinate, which explains 22% of the total variance. Together, these results reflect that CNT had a much stronger influence than graphene on soil microbial communities.

**Figure 3.**
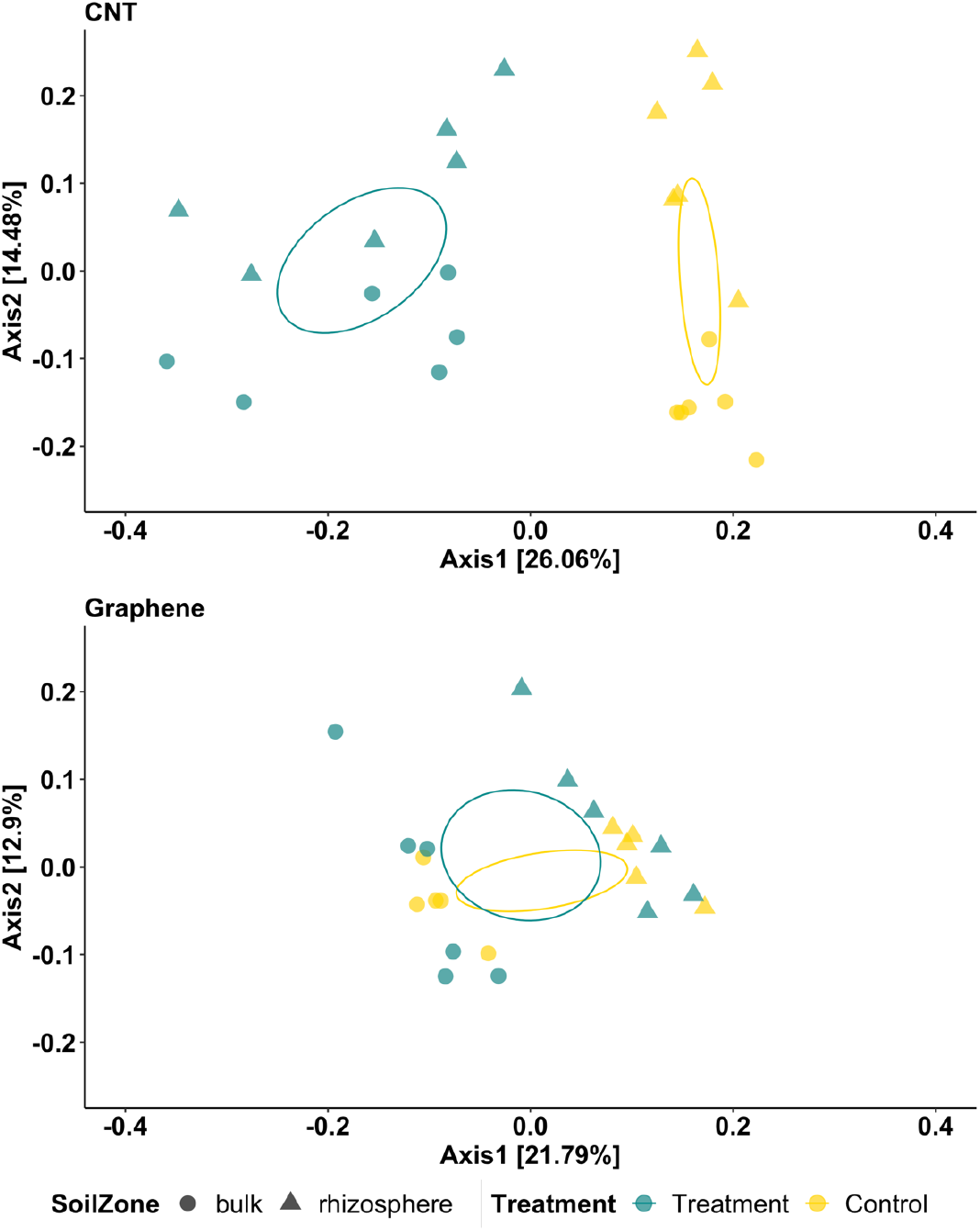
Effects of CNT (upper) and graphene (lower) on microbial beta diversity in bulk soil and the tomato rhizosphere. PCoA was based on Bray-Curtis distance of log-transformed ASV abundance. Ellipses are 95% confidence intervals for each treatment.

Studies have reported impacts of CNMs on microbial community structure in bulk soil. Previously, we reported CNT-induced shifts in microbial beta diversity in bulk soils and showed that single-walled CNT typically exerted stronger effects than multi-walled CNT, where application rate and exposure duration were influential factors.^24^ In an one-year exposure study, Ge et al. (2016) found that narrow multi-walled CNT (average diameter 7.4 nm) and graphene (average diameter 2.4 μm) significantly altered soil bacterial communities, whereas wide multi-walled CNT (average diameter >13 nm) did not.^34^ Those results were opposite to our observations here, although the multi-walled CNT and graphene used in both studies had comparable dimensions. This discrepancy may reflect the influence of experiment duration (one year versus one month) on the transformation of CNMs in the soil-plan system and thus their effects on the soil microbiome.^70^

Foremost, our results suggest that CNT influence was greater than the soil zone itself, whereas graphene did not have such influence. Similar to us, Ge et al. (2018) reported greater divergence of the soybean rhizosphere microbiome due to CNT compared to graphene, and attributed this phenomenon to the colloidal stability and differing toxicity mechanisms of these two CNMs.^35^ A study of the *S. nigrum* rhizosphere, however, found no significant effect of multi-walled CNT on either bacterial or fungal community structure.^26^ However, soils used in that study were from highly polluted historical metal mining sites, which may have introduced other confounding factors causing the discrepancy between their and our results. Alternatively, the discrepancy may be owing to rhizosphere conditions specific to the plants used (*S. nigrum* versus tomato). Research of other plant species, especially those of agricultural importance, is warranted to validate the generality of our observation.

### CNT affected more taxa in tomato-associated soil microbiomes

The top 10 bacterial phyla among all the soil samples were Acidobacteriota, Actinobacteriota, Bacteroidota, Chloroflexi, Cyanobacteria, Gemmatimonadota, Myxococcota, Planctomycetota, Proteobacteria and Verrucomicrobiota (Figure 4). As expected, many phyla had differential abundance in bulk soil versus the tomato rhizosphere. Bdellovibrionota, Elusimicrobiota, FCPU426, Fibrobacterota, Myxococcota, Nanoarchaeota, Patescibacteria, SAR324_clade(Marine_group_B), Spirochaetota, Sumerlaeota, and Verrucomicrobiota were enriched in bulk soil, whereas Abditibacteriota, Cyanobacteria, MBNT15, and Proteobacteria were enriched in the tomato rhizosphere (*p* < 0.05 in Kruskal-Wallis test).

**Figure 4.**
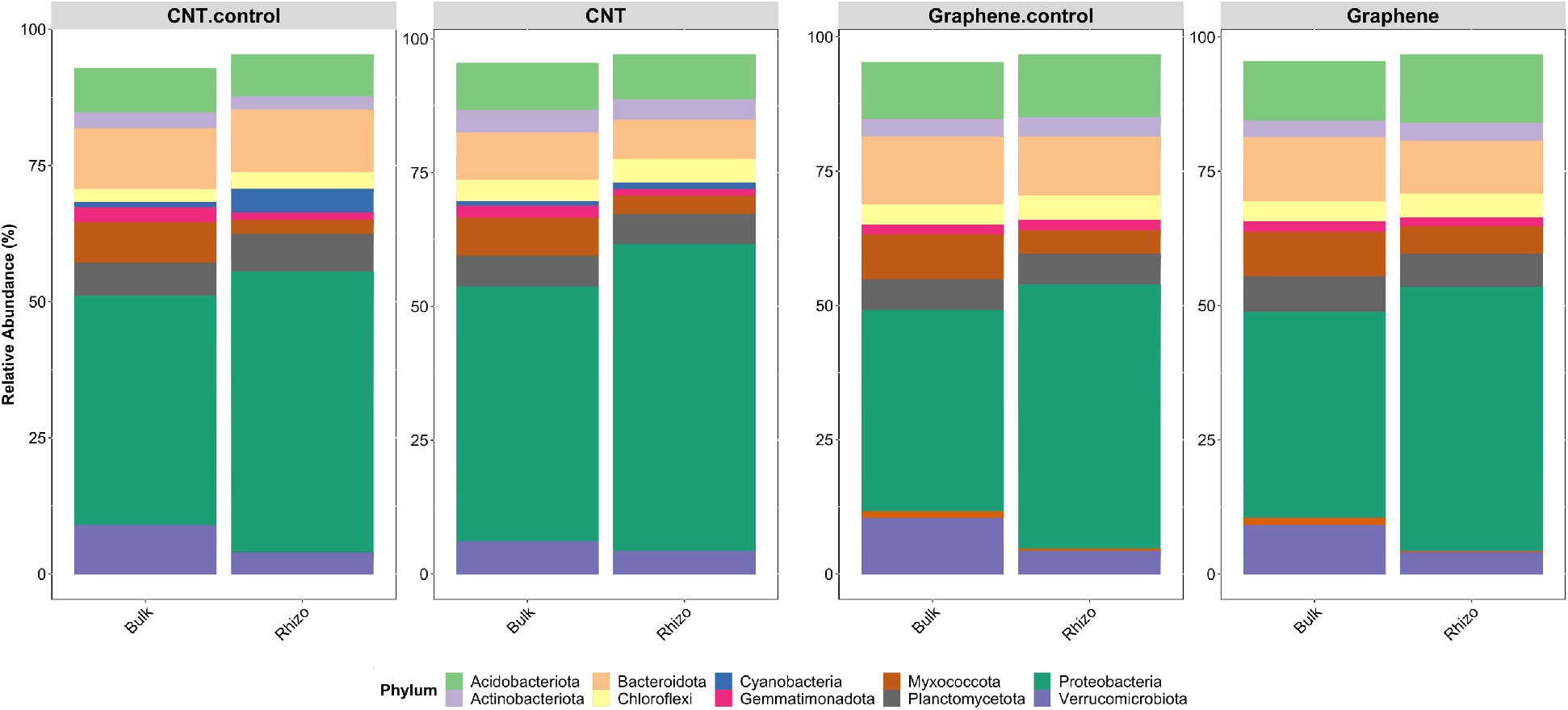
Effects of CNT (left) and graphene (right) on the relative abundance of the top 10 phyla in bulk soil and the tomato rhizosphere. Data present averages across biological replicates.

Comparing the two CNMs, CNT significantly affected more prokaryotic taxa than graphene (phylum level results in Figure 4; class level results in Figure S3). In bulk soil, CNT significantly enhanced the phylum Actinobacteriota, Chloroflexi and Sumerlaeota, but suppressed Desulfobacterota, Fibrobacterota, Firmicutes, Spirochaetota and Verrucomicrobiota, whereas graphene significantly enhanced FCPU426, Nitrospirota and Zixibacteria (*p* < 0.05 in Kruskal-Wallis test). In the tomato rhizosphere, CNT significantly enhanced Actinobacteriota and WS2, but suppressed Cyanobacteria (*p* < 0.05 in Kruskal-Wallis test), whereas graphene significantly enhanced Abditibacteriota and WPS-2 (*p* < 0.05 in Kruskal-Wallis test).

LEfSe identified a wide range of differential taxa at the phylum or class level that were most likely to explain CNT-introduced microbiome shifts (Figure 5). In bulk soil, CNT enhanced the phylum Latescibacterota and WS2, the class Blastocatellia, Subgroup 5 and Vicinamibacteria (all in Acidobacteriota phylum), Acidimicrobiia (Actinobacteriota phylum), Kapabacteria (Bacteroidota phylum), Dehalococcoidia, JG30-KF-CM66 and KD4_96 (Chloroflexi phylum), Lineage IIb (Elusimicrobiota phylum), and Chlamydiae (Verrucomicrobiota), as well as the candidate division WWE3 (Patescibacteria phylum). Taxa suppressed by CNT included the archaeal class Bathyarchaeia (Crenarchaeota phylum), the bacterial class Acidobacteriae and Holophagae (both in Acidobacteriota phylum), Coriobacteriia (Actinobacteriota phylum), Vampirivibrionia (Cyanobacteria phylum), Desulfobulbia, Desulfovibrionia, Syntrophia, Syntrophobacteria and Syntrophobacteria (all in Desulfobacterota phylum), Bacillia, Clostridia, Desulfotomaculia, Limnochordia and Negativicutes (all in Firmicutes phylum), Gemmatimonadetes (Gemmatimonadota phylum), CPR2 (Patescibacteria phylum), Planctomycetes (Planctomycetota phylum), and Spirochaetia (Spirochaetota phylum), as well as BD2-11 terrestrial group. In the rhizosphere, CNT enhanced the phylum WS2, the class Blastocatellia, Kapabacteria, KD4_96 and Thermoleophilia (all in Actinobacteriota phylum), Armatimonadia (Armatimonadota phylum), OLB14 (Chloroflexi phylum), Saccharimonadia (Patescibacteria phylum), and Alphaproteobacteria (Proteobacteria phylum). A few taxa were suppressed in the rhizosphere, including the class Vampirivibrionia, Desulfovibrionia, Clostridia, and BD2-11 terrestrial group.

**Figure 5.**
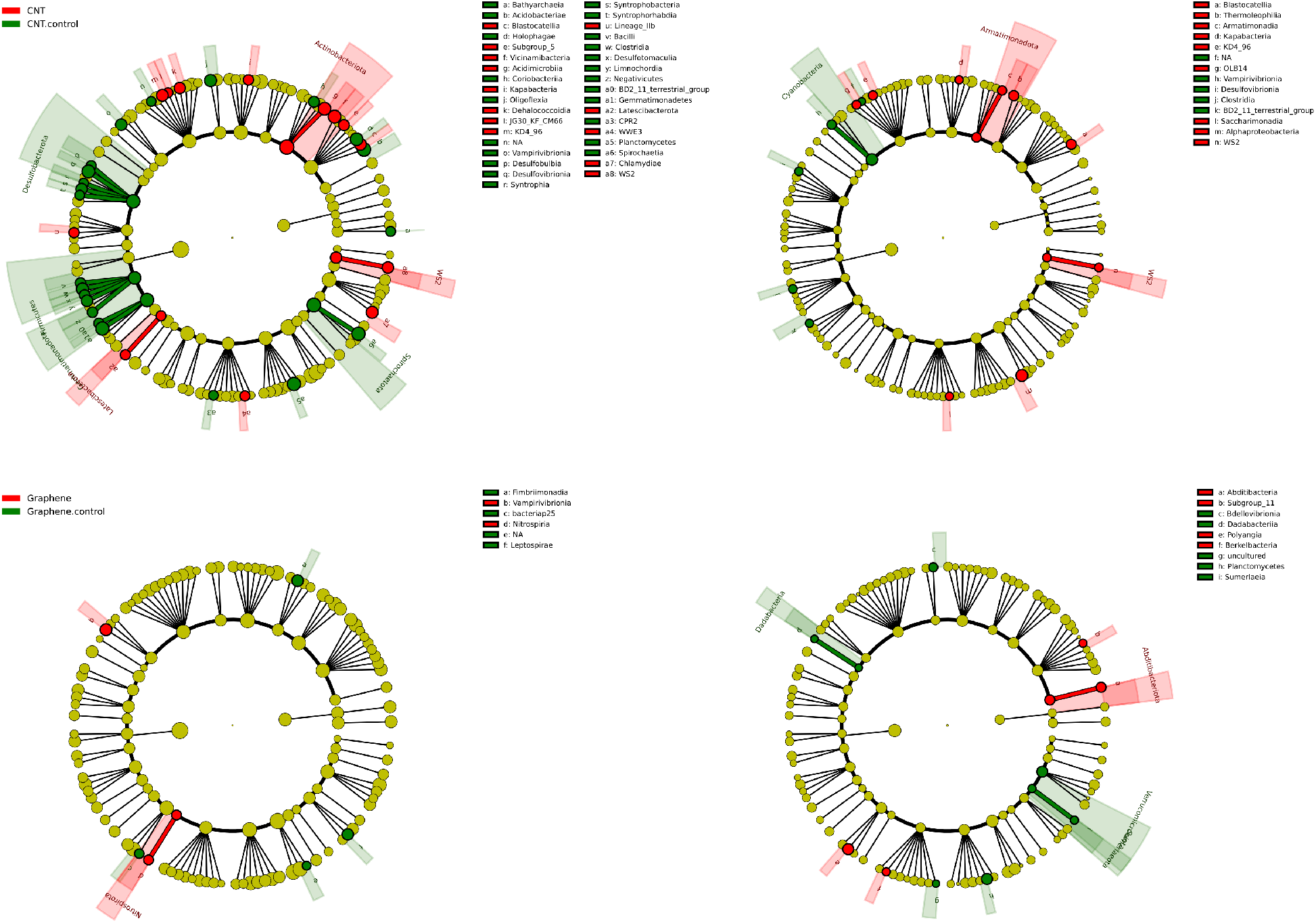
CNT influenced more microbial taxa in bulk soil (upper left) than the tomato rhizosphere (upper right). Graphene treatment influenced less microbial taxa in bulk soil (lower left) than the tomato rhizosphere (lower right). Only phyla and classes are shown here.

Compared to CNT, graphene resulted in much fewer differential taxa in bulk soil, enhancing the class Fimbriimonadia (Armatimonadota phylum), bacteriap25 (Myxococcota phylum), and Leptospirae (Spirochaetota phylum), while inhibiting the class Vampirivibrionia and Nitrospiria (both in Nitrospirota phylum) (Figure 5). In the rhizosphere, graphene increased the prevalence of the class Abditibacteria (Abditibacteriota phylum), Subgroup 11 (Acidobacteriota phylum), Polyangia (Myxococcota phylum), and Berkelbacteria (Patescibacteria phylum), but inhibited the class Bdellovibrionia (Bdellovibrionota phylum), Ddabacteriia (Dadabacteria phylum), Planctomycetes (Planctomycetota phylum), and Sumerlaeia (Sumerlaeota phylum).

In general, our results align with prior findings. In a study of the soil-tomato plant system, multi-walled CNT increased the abundance of Bacteroidota and Firmicutes but decreased Proteobacteria (particularly Alphaproteobacteria) and Verrucomicrobiota.^13^ However, that study mixed all the soil from the same pot and did not distinguish between rhizosphere and bulk soils. A bulk soil study found that single-walled CNT enhanced phyla Bacteroidota and Proteobacteria but suppressed Actinobacteriota and Chloroflexi, while multi-walled CNT increased Chloroflexi, orders Bacillales and Clostridiales under the phylum Firmicutes, and classes Deltaproteobacteria and Gammaproteobacteria under the phylum Proteobacteria.^24^ In one of the few studies of the soil-plant system, Chen et al. (2022) observed that multi-walled CNT increased Actinobacteriota and Chloroflexi but decreased Proteobacteria in the *S. nigrum* rhizosphere.^26^ Our study observed similar CNT-impacted taxa; further, we delineated the differential effects of CNT on microbial communities in the tomato rhizosphere versus surrounding bulk soil. Few studies have investigated graphene-induced microbial taxonomic changes in the plant rhizosphere. Our data indicate a much smaller influence of graphene on the soil-tomato plant system as compared to CNT. This observation explains the beta diversity trends reported earlier in this study and is consistent with others.^35^

### CNT enhanced microbial interactions in the tomato rhizosphere

Microbial interactions are essential to the structure and function of the soil microbial community.^71^ Here we observed distinct effects of CNT and graphene on microbial networks in the soil-tomato plant system. In the bulk soil, both CNMs weakened microbial interactions, illustrated by reduced network edges, particularly shorter-distance edges (Figures 6, S4, and S5). In the tomato rhizosphere, both CNMs likely strengthened microbial interactions, evident by increased edges (particularly for CNT) and/or shortened edge distance (particularly for graphene). Graph-based permutation tests further suggest that in the CNT experiment, both treatment and soil zone were significant factors shaping microbial occurrence network (*p* < 0.002 in NN tree-based permutation test), whereas in the graphene experiment, soil zone but not treatment was the significant factor shaping microbial network (*p* < 0.001 and *p* = 0.864 in NN tree-based permutation test for soil zone and treatment, respectively) (Figure S6 and S7).

**Figure 6.**
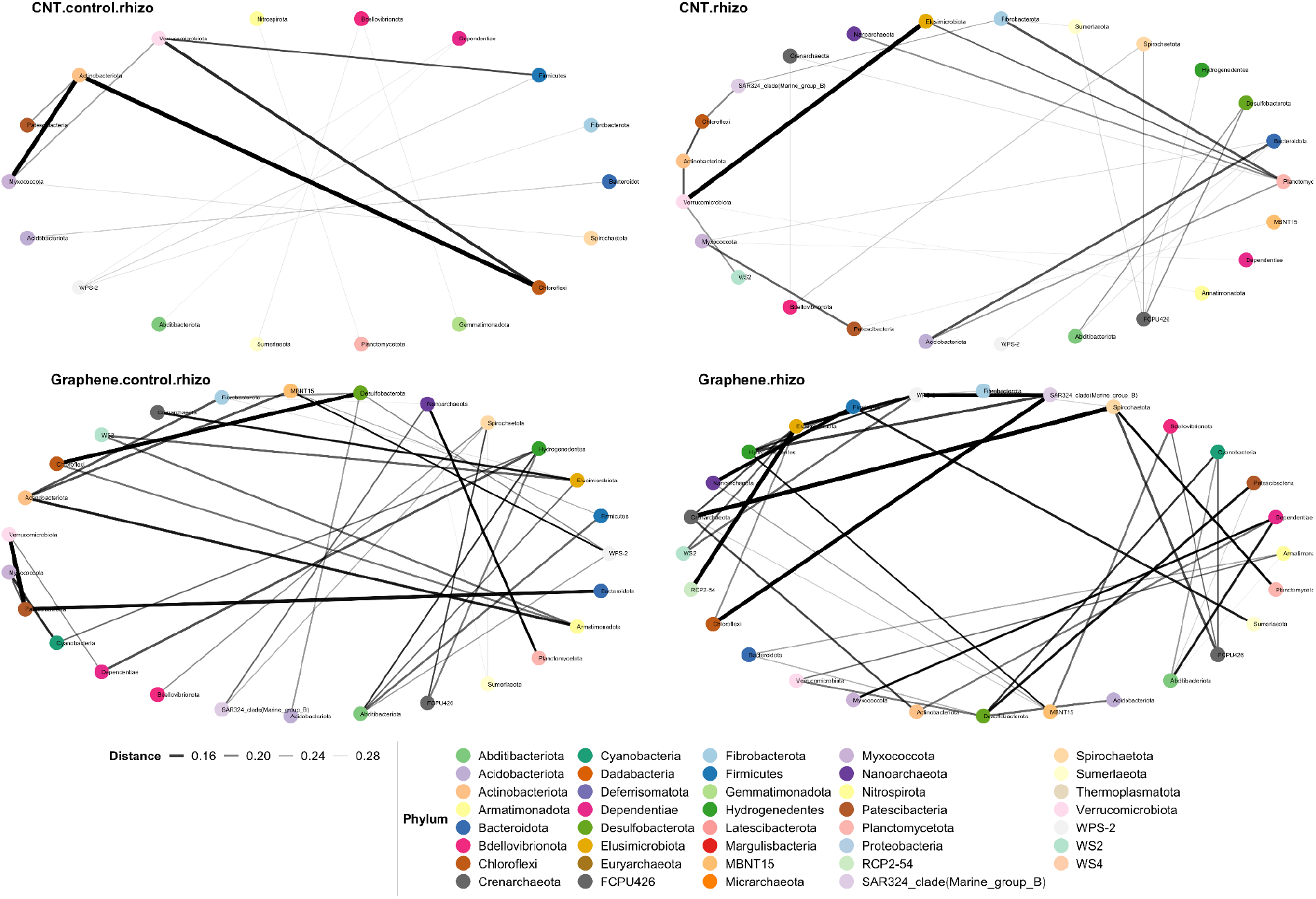
Effects of CNT and graphene on the phylum-level microbial network in the bulk soil and the tomato rhizosphere. Networks were calculated based on Bray-Curtis distance with a maximum distance of 0.3.

The generally disrupting effects of the two CNMs on the bulk soil microbiome are consistent with their suppression effects on many taxonomic groups in bulk soil as shown in Figures 4 and 5. In particular, CNT suppressed a large number of microbial taxa than graphene and thus disturbed the bulk soil microbial network more severely. On the other hand, CNT significantly increased microbial alpha diversity (Figures 1 and 2) and the abundance of various microbial taxa (Figures 4 and 5) in the tomato rhizosphere, which may contribute to strengthened microbial interactions through new microbe-microbe connections. Graphene-enhanced microbial interactions were mainly due to shorter edge distance, consistent with graphene’s limited influence on microbial alpha diversity and taxonomic shifts in the tomato rhizosphere. To our knowledge, this is the first report of CNMs enhancing microbial integrations in the plant rhizosphere. We speculate that CNMs, particularly CNT, triggered changes in the rhizosphere metabolome; in response, microbes cooperated to adapt to the CNM-modulated rhizosphere microenvironment. It should be noted, however, that correlation-based co-occurrence network does not necessarily predict ecological relationships. We envision that metabolomics analysis, integrated with metagenomics, will fully address chemical-microbe interplays in CNM-impacted plant rhizosphere.

### CNT-induced microbial functional changes

Soil microbial communities mediate critical biogeochemical processes and changes in community structure and/or composition could have a profound influence on soil function. To address CNM-induced community functional changes, we first performed 16S-based functional inference. Our results suggest that graphene did not impact microbial functions in either bulk soil or the tomato rhizosphere. In contrast, CNT affected distinct microbial pathways in bulk soil and the tomato rhizosphere (Tables S2 and S3). In bulk soil, CNT significantly impacted nitrogen metabolism (*p* < 0.001, FDR < 0.005), having negative effects on nitrogen fixation (*p* < 0.001, FDR = 0.067) (Tables S2 and S3). This consists with one of our prior studies where microarray analysis identified suppression of nitrogen fixation, nitrification, dissimilatory nitrogen reduction, and anaerobic ammonium oxidation in soils treated by multi-walled CNT at 30 or 300 mg/kg of soil.^25^ CNT-introduced downregulation of nitrogen metabolism was also seen in a *Pseudomonas aeruginosa* model strain, where 10 mg/L multi-walled CNT induced differential transcriptional regulation of 2.5 times more genes than graphene at the same concentration.^72^

In the rhizosphere, CNT significantly impacted the pathways of degradation of aromatic compounds, steroid degradation, phenylalanine metabolism, and methane metabolism, negatively affecting the modules of benzene degradation, anaerobic toluene degradation, and tocopherol biosynthesis (*p* < 0.005, FDR < 0.05) (Tables S2 and S3). Interestingly, CNT affected F_420_ biosynthesis both positively (gene *cofD* and *cofE*) and negatively (gene *cofG* and *cofH*) (Figure S8). Prior research of CNM impact on the functionality of plant rhizosphere microbiomes focused on legumes and showed that CNMs may impair symbiotic nitrogen fixation by inhibiting the initiation of nodulation or symbiosis formation.^35,73^ We did not observe significant impact of either CNT or graphene on microbial nitrogen metabolism in the tomato rhizosphere, despite of their negative impact on microbial nitrogen fixation in bulk soil. Little is known about CNM-induced microbial functional shifts in the rhizosphere of non-leguminous plants. Here the negative effects of CNT on microbial traits might be attributed to the reduced abundance of the corresponding functional groups. For example, tocopherols (lipophilic antioxidants within the vitamin-E family) are synthesized exclusively by photosynthetic organisms, including some cyanobacteria.^74^ Our data showed that CNT significantly inhibited Cyanobacteria and thus negatively affected tocopherol biosynthesis in the tomato rhizosphere. The influence of CNT on F_420_ biosynthesis has not been reported previously. Cofactor F_420_, historically known as a methanogenic redox cofactor, is widely distributed across the bacterial and archaeal domains, catalyzing challenging redox reactions including key steps in methanogenesis, xenobiotic degradation, and antibiotic biosynthesis. Prokaryotic F_420_ synthesize pathways have three variants, existing in Euryarchaeota (*cofD* and *cofE* present), Actinobacteriota and Chloroflexi, and Proteobacteria (*cofE* present), but all start with deazaflavin Fo synthesis mediated by *cofG/H* or *fbiC*.^75^ Our data suggest that CNT increased the phylum Actinobacteriota, the class Alphaproteobacteria and members of Chloroflexi, but decreased the class Clostridia in the tomato rhizosphere. These groups are known F_420_ producers and their abundance changes might affect F_420_ biosynthesis pathways.

To further evaluate CNM effects on the functionality of soil microbial communities, we employed EcoPlate assay, a culture-dependent analysis of community-level substrate metabolism. Due to the limited amount of rhizosphere soils, only bulk soils were analyzed. Out of the 31 tested substrates, twenty seven and sixteen substrates were less used in soils treated by CNT or graphene, respectively, in comparison to control soils (Figure 8). CNT led to less use of one amine, two amino acids, one carbohydrate, two carboxylic acids and one polymer, whereas graphene led to less use of one amino acid and one carboxylic acid (*p* < 0.05 in Kruskal-Wallis test). Specifically, CNT reduced microbial use of labile substrates (amines/amid, amino acids), whereas graphene promoted microbial use of both labile (amines/amid, amino acids) and complex substrates (polymers). Interestingly, CNT seemed to promote basal respiration, whereas graphene did not have such an effect (Figure S10). Ge et al. (2016) also found higher soil basal respiration in CNT-treated soils than controls after one month of exposure, although over one year, there was no significant difference across treatments.^34^ Our study maintained a higher water content than Ge et al. (2016) (11% versus 5%), which may have contributed to greater soil microbial activities by facilitating the diffusion of soluble substrates while not limiting the diffusion of oxygen.^76,77^ Together, EcoPlate assay and basal respiration assessment suggest that CNT treatment narrowed the substrate spectrum of microbes but increased their activity in the bulk soil. Whether similar phenomena also occur in the plant rhizosphere requires further investigation. Specifical attention should be paid to the microbial metabolism of root exudates which are rich in amino acids, organic acids, sterols, and others.^78^

**Figure 8.**
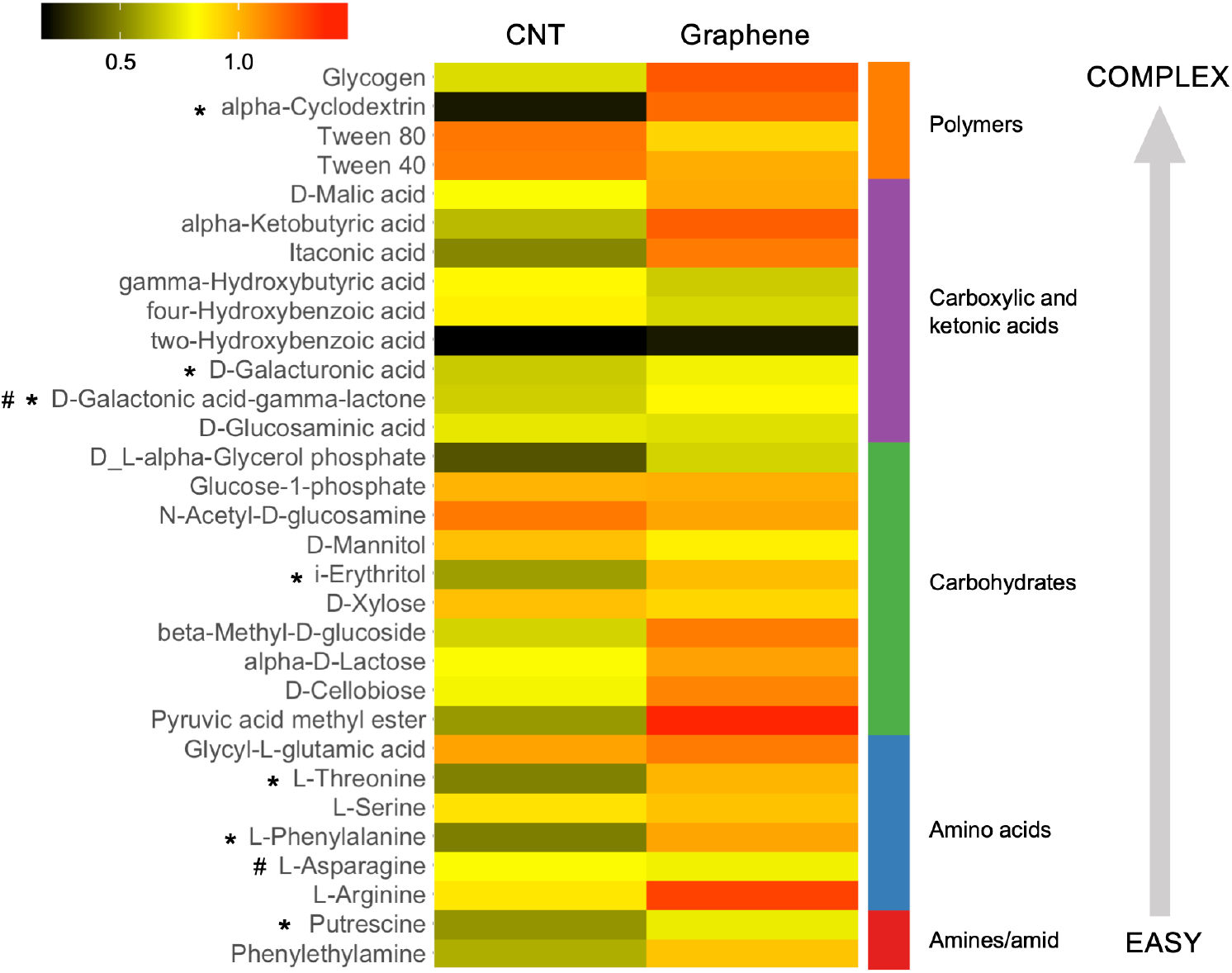
Heatmap showing CNT and graphene effects on microbial substrate utilization. AWCD values were averaged across technique replicates and pot replicates and normalized against corresponding controls. The color key indicates fold change. Asterisks and number signs indicate substrate compound significantly affected by CNT or graphene, respectively, as compared to controls (*p* < 0.05 in Kruskal-Wallis test). Substrate categories are presented on the right side of the heatmap with substrate complexity increasing from bottom to top as annotated by the grey arrow.

## Conclusion

To realize the full potential of CNMs for sustainable crop production, it is imperative to elucidate how CNMs influence the plant holobiont which comprises the host plant and its microbiota. This study depict the effects of two widely used CNMs, multi-walled CNT and graphene, on microbiomes in the plant ectorhizosphere and surrounding bulk soil. We focused on tomato, one of the most consumed vegetable crops around the world, and identified greater effects of CNT than graphene on the tomato ectorhizosphere microbiome. CNT significantly shaped microbial diversity, community composition, microbe-microbe interaction, and likely also metabolic activities. Foremost, compared to their counterparts in bulk soil, microbiomes in the tomato ectorhizosphere displayed stronger and/or unique shifts in response to CNT, including greater community divergence, strengthened microbial interaction, and potentially distinct functional shifts. The significance of these results is twofold. First, it highlights the differential modulation effects of CNT and graphene on plant rhizosphere microbiomes and the potential involvement of root exudates that are themselves under CNM modulation. Second, it emphasizes the need of addressing the influence of rhizosphere microbiome alterations on plant fitness. From a holobiont perspective, root extrudates play an important role in the recruitment and assembly of the host plant’s rhizosphere microbiome, which in turn confers fitness advantages (e.g., nutrient update, stress tolerance, pathogen resistance) to the plant host. Further research on the interplays between roots and microbes in the CNM-impacted rhizosphere will help unveil mechanisms underlying CNM-introduced plant phenotypical changes. It will also shed light on CNM-mediated manipulation of plant rhizosphere microbiomes for more sustainable crop production.

## Acknowledgments

Y. You acknowledges support from Idaho State University (the Department of Biological Science, the College of Science and Engineering) and the State University of New York College of Environmental Science and Forestry. Y. You also acknowledges Oak Ridge Associated Universities to sponsor the seminar series “From Genome to Exposome: State of Biosystem Signaling in Response to Emerging Environmental Stressors” that inspired this research. P. Kerner was supported by the Center for Ecological Research and Education at Idaho State University. We thank Dr. Kathleen Lohse (Idaho State University) for providing EGM-4 gas analyzer, and Lisa McDougall and Jason Werth (Molecular Research Core Facility, Idaho State University) for completing Illumina sequencing while facing all the challenges associated with the COVID-19 pandemic. M. Khodakovskaya acknowledges funding from USDA-NIFA (AFRI 2017-07886) and NASA-EPSCoR (NNH16ZHA0055C) which allowed to creation of infrastructure for experimental work with nanomaterials in UA Little Rock.

## References

1. S.M. Rodrigues, P. Demokritou, N. Dokoozlian, C.O. Hendren, B. Karn, M.S. Mauter, O.A. Sadik, M. Safarpour, J.M. Unrine, J. Viers and P. Welle, Nanotechnology for sustainable food production: promising opportunities and scientific challenges, Environ. Sci.: Nano., 2017, 4, 767–81.

2. I.O. Adisa, V.L.R. Pullagurala, J.R. Peralta-Videa, C.O. Dimkpa, W.H. Elmer, J.L. Gardea-Torresdey and J.C. White, Recent advances in nano-enabled fertilizers and pesticides: a critical review of mechanisms of action, Environ. Sci.: Nano., 2019, 6, 2002–2030.

3. National Academies of Sciences, Engineering, and Medicine, Science Breakthroughs to Advance Food and Agricultural Research by 2030. National Academies Press, 2019.

4. M. Usman, M. Farooq, A. Wakeel, A. Nawaz, S.A. Cheema, H. ur Rehman, I. Ashraf and M. Sanaullah, Nanotechnology in agriculture: Current status, challenges and future opportunities, Sci. Total Environ., 2020, 721, 137778.

5. A. Gogos, K. Knauer and T.D. Bucheli, Nanomaterials in plant protection and fertilization: current state, foreseen applications, and research priorities, J. Agric. Food Chem., 2012, 60, 9781–9792.

6. P. Wang, E. Lombi, F.J. Zhao and P.M. Kopittke, Nanotechnology: a new opportunity in plant sciences, Trends Plant Sci., 2016, 21, 699–712.

7. T. Hofmann, G.V. Lowry, S. Ghoshal, N. Tufenkji, D. Brambilla, J.R. Dutcher, L.M. Gilbertson, J.P. Giraldo, J.M. Kinsella, M.P. Landry and W. Lovell, Technology readiness and overcoming barriers to sustainably implement nanotechnology-enabled plant agriculture, Nat. Food, 2020, 1, 416–425.

8. H. Yang, C. Wang, F. Chen, L. Yue, X. Cao, J. Li, X. Zhao, F. Wu, Z. Wang and B. Xing, Foliar carbon dot amendment modulates carbohydrate metabolism, rhizospheric properties and drought tolerance in maize seedling, Sci. Total Environ., 2022, 809, 151105.

9. S.K. Verma, A.K. Das, S. Gantait, V. Kumar and E. Gurel, Applications of carbon nanomaterials in the plant system: A perspective view on the pros and cons. Sci. Total Environ., 2019, 667, 485–499.

10. A. Mukherjee, S. Majumdar, A.D. Servin, L. Pagano, O.P. Dhankher and J.C. White, Carbon nanomaterials in agriculture: a critical review, Front. Plant Sci., 2016, 7, 172.

11. M.H. Lahiani, E. Dervishi, J. Chen, Z. Nima, A. Gaume, A.S. Biris and M.V. Khodakovskaya, Impact of carbon nanotube exposure to seeds of valuable crops, ACS Appl. Mater. Interfaces, 2013, 5, 7965–7973.

12. M.V. Khodakovskaya, K. de Silva, D.A. Nedosekin, E. Dervishi, A.S. Biris, E.V. Shashkov, E.I. Galanzha and V.P. Zharov, Complex genetic, photothermal, and photoacoustic analysis of nanoparticle-plant interactions, Proc. Natl. Acad. Sci. U. S. A., 2011, 108, 1028–1033.

13. M.V. Khodakovskaya, B.S. Kim, J.N. Kim, M. Alimohammadi, E. Dervishi, T. Mustafa and C.E. Cernigla, Carbon nanotubes as plant growth regulators: effects on tomato growth, reproductive system, and soil microbial community, Small, 2013, 9, 115–123.

14. M.H. Lahiani, E. Dervishi, I. Ivanov, J. Chen and M. Khodakovskaya, Comparative study of plant responses to carbon-based nanomaterials with different morphologies, Nanotechnology, 2016, 27, 265102.

15. M.H. Lahiani, Z.A. Nima, H. Villagarcia, A.S. Biris and M.V. Khodakovskaya, Assessment of effects of the long-term exposure of agricultural crops to carbon nanotubes, J. Agric. Food Chem., 2017, 66, 6654–6662.

16. D.L. McGehee, M. Alimohammadi and M.V. Khodakovskaya, Carbon-based nanomaterials as stimulators of production of pharmaceutically active alkaloids in cell culture of Catharanthus roseus, Nanotechnology, 2019, 30, 275102.

17. K. Pandey, M.H. Lahiani, V.K. Hicks, M.K. Hudson, M.J. Green and M. Khodakovskaya, Effects of carbon-based nanomaterials on seed germination, biomass accumulation and salt stress response of bioenergy crops, PloS ONE, 2018, 13, e0202274.

18. S. Rezaei Cherati, S. Shanmugam, K. Pandey and M.V. Khodakovskaya, Whole-transcriptome responses to environmental stresses in agricultural crops treated with carbon-based nanomaterials, ACS Appl. Bio Mater., 2021, 4, 4292–4301.

19. M.V. Khodakovskaya, K. de Silva, D.A. Nedosekin, E. Dervishi, A.S. Biris, E.V. Shashkov, E.I. Galanzha and V.P. Zharov, Complex genetic, photothermal, and photoacoustic analysis of nanoparticle-plant interactions, Proc. Natl. Acad. Sci. U. S. A., 2011, 108, 1028–1033.

20. Z. Cao, H. Zhou, L. Kong, L. Li, R. Wang and W. Shen, A novel mechanism underlying multi-walled carbon nanotube-triggered tomato lateral root formation: the involvement of nitric oxide, Nanoscale Res. Lett., 2020, 15, 1–10.

21. M.H. Lahiani, J. Chen, F. Irin, A.A. Puretzky, M.J. Green and M.V. Khodakovskaya, Interaction of carbon nanohorns with plants: uptake and biological effects, Carbon, 2015, 81, 607–619.

22. Z. Chen, J. Zhao, J. Song, S. Han, Y. Du, Y. Qiao, Z. Liu, J. Qiao, W. Li, J. Li and H. Wang, Influence of graphene on the multiple metabolic pathways of Zea mays roots based on transcriptome analysis, PLoS One, 2021, 16, e0244856.

23. S. Rezaei Cherati, M. Anas, S. Liu, S. Shanmugam, K. Pandey, S. Angtuaco, R. Shelton, A.N. Khalfaoui, S.V. Alena, E. Porter and T. Fite, Comprehensive Risk Assessment of Carbon Nanotubes Used for Agricultural Applications, ACS Nano, 2022, 16, 12061–12072.

24. F. Wu, Y. You, X. Zhang, H. Zhang, W. Chen, Y. Yang, D. Werner, S. Tao and X. Wang, Effects of various carbon nanotubes on soil bacterial community composition and structure, Environ. Sci. Technol., 2019, 53, 5707–5716.

25. F. Wu, Y. You, D. Werner, S. Jiao, J. Hu, X. Zhang, Y. Wan, J. Liu, B. Wang and X. Wang, Carbon nanomaterials affect carbon cycle-related functions of the soil microbial community and the coupling of nutrient cycles, J. Hazard. Mater., 2020, 390, 122144.

26. X. Chen, J. Wang, Y. You, R. Wang, S. Chu, Y. Chi, K. Hayat, N. Hui, X. Liu, D. Zhang and P. Zhou, When nanoparticle and microbes meet: The effect of multi-walled carbon nanotubes on microbial community and nutrient cycling in hyperaccumulator system, J. Hazard. Mater., 2022, 423, 126947.

27. R. Mendes, M. Kruijt, I. De Bruijn, E. Dekkers, M. Van Der Voort, J.H. Schneider, Y.M. Piceno, T.Z. DeSantis, G.L. Andersen, P.A. Bakker and J.M. Raaijmakers, Deciphering the rhizosphere microbiome for disease-suppressive bacteria, Science, 2011, 332, 1097–1100.

28. A.H. Ahkami, R.A. White Iii, P.P. Handakumbura and C Jansson, Rhizosphere engineering: enhancing sustainable plant ecosystem productivity, Rhizosphere, 2017, 3, 233–243.

29. M.J. Kwak, H.G. Kong, K. Choi, S.K. Kwon, J.Y. Song, J. Lee, P.A. Lee, S.Y. Choi, M. Seo, H.J. Lee and E.J. Jung, Rhizosphere microbiome structure alters to enable wilt resistance in tomato, Nat. Biotechnol., 2018, 36, 1100–1109.

30. P. Trivedi, J.E. Leach, S.G. Tringe, T. Sa and B.K. Singh, Plant–microbiome interactions: from community assembly to plant health, Nat. Rev. Microbiol., 2020, 18, 607–621.

31. Q. Qu, Z. Zhang, W.J.G.M. Peijnenburg, W. Liu, T. Lu, B. Hu, J. Chen, J. Chen, Z. Lin and H. Qian, Rhizosphere microbiome assembly and its impact on plant growth, J. Agric. Food Chem., 2020, 68, 5024–5038.

32. P. Vandenkoornhuyse, A. Quaiser, M. Duhamel, A. Le Van and A. Dufresne, The importance of the microbiome of the plant holobiont, New Phytol., 2015, 206, 1196–1206.

33. Y. You, K.K. Das, H. Guo, C.W. Chang, M. Navas-Moreno, J.W. Chan, P. Verburg, S.R. Poulson, X. Wang, B. Xing and Y. Yang, Microbial transformation of multiwalled carbon nanotubes by Mycobacterium vanbaalenii PYR-1, Environ. Sci. Technol., 2017, 51, 2068–2076.

34. Y. Ge, J.H. Priester, M. Mortimer, C.H. Chang, Z. Ji, J.P. Schimel and P.A. Holden, Long-term effects of multiwalled carbon nanotubes and graphene on microbial communities in dry soil, Environ. Sci. Technol., 2016, 50, 3965–3974.

35. Y. Ge, C. Shen, Y. Wang, Y.Q. Sun, J.P. Schimel, J.L. Gardea-Torresdey and P.A. Holden, Carbonaceous nanomaterials have higher effects on soybean rhizosphere prokaryotic communities during the reproductive growth phase than during vegetative growth, Environ. Sci. Technol., 2018, 52, 6636–6646.

36. F. Zhao, X. Xin, Y. Cao, D. Su, P. Ji, Z. Zhu and Z. He, Use of carbon nanoparticles to improve soil fertility, crop growth and nutrient uptake by corn (Zea mays L.), Nanomaterials, 2021, 11, 2717.

37. F. Gottschalk, T. Sun and B. Nowack, Environmental concentrations of engineered nanomaterials: review of modeling and analytical studies, Environ. Pollut., 2013, 181, 287–300.

38. A.A. Keller and A. Lazareva, Predicted releases of engineered nanomaterials: From global to regional to local, Environ. Sci. Technol., 2014, 1, 65–70.

39. M.R. McPherson, P. Wang, E.L. Marsh, R.B. Mitchell and D.P. Schachtman, Isolation and analysis of microbial communities in soil, rhizosphere, and roots in perennial grass experiments, J. Vis. Exp., 2018, 137, 57932.

40. T. Simmons, D.F. Caddell, S. Deng and D. Coleman-Derr, Exploring the root microbiome: extracting bacterial community data from the soil, rhizosphere, and root endosphere, J. Vis. Exp., 2018, 135, 57561.

41. A. Gryta, M. Frąc and K. Oszust, 2014. The application of the Biolog EcoPlate approach in ecotoxicological evaluation of dairy sewage sludge, Appl. Biochem. Biotechnol., 174, 1434–1443.

42. M.R. Green and J. Sambrook, 2016. Precipitation of DNA with ethanol. Cold Spring Harbor Protocols, 2016, 12, pdb-prot093377.

43. Y. You, K. Aho, K.A. Lohse, S.G. Schwabedissen, R.N. Ledbetter and T.S. Magnuson, Biological soil crust bacterial communities vary along climatic and shrub cover gradients within a sagebrush steppe ecosystem, Front. Microbiol., 2021, 12, 1096.

44. J.G. Caporaso, C.L. Lauber, W.A. Walters, D. Berg-Lyons, J. Huntley, N. Fierer, S.M. Owens, J. Betley, L. Fraser, M. Bauer and N. Gormley, Ultra-high-throughput microbial community analysis on the Illumina HiSeq and MiSeq platforms, The ISME J., 2012, 6, 1621–1624.

45. C. Marotz, A. Sharma, G. Humphrey, N. Gottel, C. Daum, J.A. Gilbert, E. Eloe-Fadrosh and R. Knight, Triplicate PCR reactions for 16S rRNA gene amplicon sequencing are unnecessary, BioTechniques, 2019, 67, 29–32.

46. A. Apprill, S. McNally, R. Parsons and L. Weber, Minor revision to V4 region SSU rRNA 806R gene primer greatly increases detection of SAR11 bacterioplankton, Aquat. Microb. Ecol., 2015, 75, 129–137.

47. A.E. Parada, D.M. Needham and J.A. Fuhrman, Every base matters: Assessing small subunit rRNA primers for marine microbiomes with mock communities, time series and global field samples, Environ. Microbiol., 2016, 18, 1403–1414.

48. W. Walters, E.R. Hyde, D. Berg-Lyons, G. Ackermann, G. Humphrey, A. Parada, J.A. Gilbert, J.K. Jansson, J.G. Caporaso, J.A. Fuhrman and A. Apprill, Improved bacterial 16S rRNA gene (V4 and V4-5) and fungal internal transcribed spacer marker gene primers for microbial community surveys, mSystems, 2016, 1, e00009–15.

49. S. Andrews, 2010. FastQC: a quality control tool for high throughput sequence data, http://www.bioinformatics.babraham.ac.uk/projects/fastqc.

50. E. Bolyen, J.R. Rideout, M.R. Dillon, N.A. Bokulich, C.C. Abnet, G.A. Al-Ghalith, H. Alexander, E.J. Alm, M. Arumugam, F. Asnicar and Y. Bai, Reproducible, interactive, scalable and extensible microbiome data science using QIIME 2, Nat. Biotechnol., 2019, 37, 852–857.

51. B.J. Callahan, P.J. McMurdie, M.J. Rosen, A.W. Han, A.J.A. Johnson and S.P. Holmes, DADA2: high-resolution sample inference from Illumina amplicon data, Nat. Methods, 2016, 13, 581–583.

52. B.J. Callahan, P.J. McMurdie and S.P. Holmes, Exact sequence variants should replace operational taxonomic units in marker-gene data analysis, The ISME J., 2017, 11, 2639–2643.

53. C. Quast, E. Pruesse, P. Yilmaz, J. Gerken, T. Schweer, P. Yarza, J. Peplies and F.O. Glöckner, The SILVA ribosomal RNA gene database project: improved data processing and web-based tools, Nucleic Acids Res., 2012, 41, D590–D596.

54. N.A. Bokulich, B.D. Kaehler, J.R. Rideout, M. Dillon, E. Bolyen, R. Knight, G.A. Huttley and J.G. Caporaso, Optimizing taxonomic classification of marker-gene amplicon sequences with QIIME 2’s q2-feature-classifier plugin, Microbiome, 2018, 6, 1–17.

55. C. Lozupone and R. Knight, UniFrac: a new phylogenetic method for comparing microbial communities, Appl. Environ. Microbiol., 2005, 71, 8228–8235.

56. N. Segata, J. Izard, L. Waldron, D. Gevers, L. Miropolsky, W.S. Garrett and C. Huttenhower, Metagenomic biomarker discovery and explanation, Genome Biol., 2011, 12, 1–18.

57. G.M. Douglas, V.J. Maffei, J.R. Zaneveld, S.N. Yurgel, J.R. Brown, C.M. Taylor, C. Huttenhower and M.G. Langille, PICRUSt2 for prediction of metagenome functions, Nat. Biotechnol., 2020, 38, 685–688.

58. M.I. Love, W. Huber and S. Anders, Moderated estimation of fold change and dispersion for RNA-seq data with DESeq2, Genome Biol., 2014, 15, 1–21.

59. J. Chong, P. Liu, G. Zhou and J. Xia, Using MicrobiomeAnalyst for comprehensive statistical, functional, and meta-analysis of microbiome data, Nat. Protoc., 2020, 15, 799–821.

60. R.C. Team, 2013. R: A language and environment for statistical computing.

61. J. Oksanen, F.G. Blanchet, R. Kindt, P. Legendre, P.R. Minchin, R.B. O’hara, G.L. Simpson, P. Solymos, M.H.H. Stevens, H. Wagner and M.J. Oksanen, 2013. Package ‘vegan’, Community Ecology Package, version 2(9), 1–295.

62. P.J. McMurdie and S. Holmes, phyloseq: an R package for reproducible interactive analysis and graphics of microbiome census data, PloS One, 2013, 8, e61217.

63. H. Wickham, ggplot2, Wiley Interdisciplinary Reviews: Computational Statistics, 2011, 3, 180–185.

64. H. Wickham, The tidyverse, R package version, 2017, 1, 836.

65. P.L. Buttigieg and A. Ramette, A guide to statistical analysis in microbial ecology: a community-focused, living review of multivariate data analyses, FEMS Microbiol. Ecol., 2014, 90, 543–550.

66. M.J. Anderson, A new method for non-parametric multivariate analysis of variance, Austral Ecol., 2001, 26, 32–46.

67. B.J. Callahan, K. Sankaran, J.A. Fukuyama, P.J. McMurdie and S.P. Holmes, Bioconductor workflow for microbiome data analysis: from raw reads to community analyses, F1000Research, 2016, 5.

68. Y. Wang, C.H. Chang, Z. Ji, D.C. Bouchard, R.M. Nisbet, J.P. Schimel, J.L. Gardea-Torresdey and P.A. Holden, Agglomeration determines effects of carbonaceous nanomaterials on soybean nodulation, dinitrogen fixation potential, and growth in soil, ACS Nano, 2017, 11, 5753–5765.

69. R. Baby, B. Saifullah and M.Z. Hussein, Carbon nanomaterials for the treatment of heavy metal-contaminated water and environmental remediation, Nanoscale Res. Lett., 2019, 14, 1–17.

70. P. Zhang, Z. Guo, Z. Zhang, H. Fu, J.C. White and I. Lynch, Nanomaterial transformation in the soil–plant system: Implications for food safety and application in agriculture, Small, 2020, 16, 2000705.

71. J.K. Jansson and K.S. Hofmockel, The soil microbiome—from metagenomics to metaphenomics, Curr. Opin. Microbiol., 2018, 43, 162–168.

72. M. Mortimer, N. Devarajan, D. Li and P.A. Holden, Multiwall carbon nanotubes induce more pronounced transcriptomic responses in Pseudomonas aeruginosa PG201 than graphene, exfoliated boron nitride, or carbon black, ACS Nano, 2018, 12, 2728–2740.

73. M. Mortimer, D. Li, Y. Wang and P.A. Holden, Physical properties of carbon nanomaterials and nanoceria affect pathways important to the nodulation competitiveness of the symbiotic N_2_-fixing bacterium Bradyrhizobium diazoefficiens, Small, 2020, 16, 1906055.

74. V.I. Lushchak and N.M. Semchuk, Tocopherol biosynthesis: chemistry, regulation and effects of environmental factors. Acta Physiol. Plant, 2012, 34, 1607–1628.

75. R. Grinter and C. Greening, Cofactor F420: An expanded view of its distribution, biosynthesis and roles in bacteria and archaea, FEMS Microbiol. Rev., 2021, 45, fuab021.

76. E.A. Davidson, I.A. Janssens and Y. Luo, On the variability of respiration in terrestrial ecosystems: moving beyond Q10, Glob. Change Biol., 2006, 12, 154–164.

77. M. Xu and H. Shang, Contribution of soil respiration to the global carbon equation, J. Plant Physiol., 2016, 203, 16–28.

78. V. Vives-Peris, C. de Ollas, A. Gómez-Cadenas and R.M. Pérez-Clemente, Root exudates: from plant to rhizosphere and beyond, Plant Cell Rep., 2020, 39, 3–17.

