## Supplementary Information for "Differential effects of carbon nanotube and graphene on the tomato rhizosphere microbiome"

Number of Pages: 23
Number of Figures: 9
Number of Tables: 3

### Materials and Methods

#### *Soil physicochemical property analyses*

Tomato plants were grown in soils receiving either carbon nanotube (CNT) or graphene treatment, with appropriate controls included in each exposure experiment (5-6 biological replicates (5 for graphene and 6 for CNT) x 2 conditions (treatment or control) x 2 carbon nanomaterials (graphene or CNT) = 22 pots in total). After the exposure experiment, tomato plants were carefully removed. Bulk soil was mixed within each of the 22 experimental pots.

For each carbon nanomaterial and experimental condition, bulk soils from three replicate pots were mixed in equal amounts and subjected to soil property analysis at the Environmental Analytical Laboratory, Brigham Young University (Provo, UT) using established protocols (https://pws.byu.edu/eal). Briefly, pH and electrical conductivity (EC) were measured using meters on a saturated soil paste. Ammonium ($NH_4$-N) and nitrate ($NO_3$-N) were extracted with 2 M potassium chloride (KCl) and measured on a Rapid Flow Analyzer (Quick Chem 8500, Lachat Instruments, Loveland, Colorado, USA). Cation exchange capacity (CEC) was measured with the ammonium replacement method on the same instrument. Organic matter (OM; in %) was measured by dichromate oxidation. Total C (%) and N (%) were determined based on combustion on an element analyzer (TruSpec CN Determinator, LECO Instruments, MI, USA). Phosphorus (P) was extracted with 0.5 M sodium bicarbonate ($NaHCO_3$) according to Olsen's method.[1] Exchangeable potassium (K) was extracted with ammonium acetate, sulfate ($SO_4$-S) was extracted with monocalcium phosphate, and micronutrients zinc (Zn), iron (Fe), manganese (Mn) and copper (Cu) were extracted with diethylenetriamine pentaacetate (DTPA). The extracted analytes were measured by inductively coupled plasma optical emission spectrometry (ICP-OES) (iCAP 7400, Thermo Electron, WI, USA). Calcium carbonate ($CaCO_3$) was measured according to Allison et al. (1965).[2]

#### *Soil basal respiration assessment*

To evaluate CNM effects on the functionality of the soil microbial community, we first performed EcoPlate assay to compare substrate utilization patterns in bulk soils harvested at the conclusion of the exposure experiment (detailed in the main text). Next we used a microcosm setup to measure soil basal respiration (Figure S1). Bulk soil collected from a pot was thoroughly mixed and 10 grams were transferred to a sterile 50 mL amber serum bottle. The water content was maintained at 11% (w/w), a typical field capacity, using sterile Milli-Q water. Each serum bottle was sealed with a rubber septum and aluminum crimp, and incubated for 1 hour in a growth chamber (model A1000, Conviron, Winnipeg, Canada) at 25 °C and under

50% humidity. Gas accumulated in bottle headspace was sampled using a 30 mL polypropylene syringe and immediately injected into an EGM-4 gas analyzer (PP systems, Amesbury, MA) for $CO_2$ measurement. Ambient $CO_2$ level was measured before each sample to ensure analytical consistency.

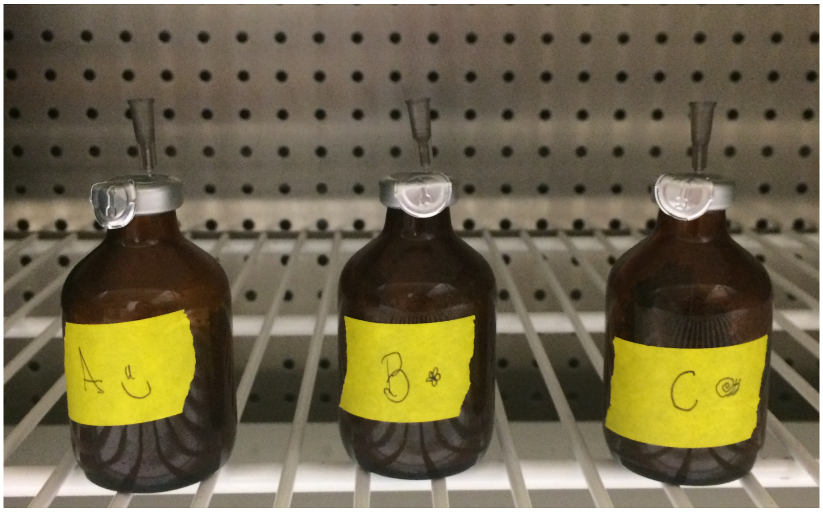


**Figure S1.** Serum bottle microcosm for bulk soil respiration potential assessment. Bottles were incubated for 1 hour in a controlled growth chamber.

### Results and Discussion

***Changes in soil property due to carbonaceous nanomaterials (CNMs)***

It should be noted that the CNT and graphene experiments were conducted using two different soils. Comparisons between bulk soils harvested from the control and treatment pots showed CNT and graphene treatment resulted in differential changes in soil property (Table S1).

**Table S1.** Properties of bulk soils from the control and treatment pots. Greatest relative changes (>=20%) were highlighted in bold.

| | CNT.control | CNT | Relative change | Graphene.control | Graphene | Relative change |
|---|---|---|---|---|---|---|
| pH | 6.66 | 6.54 | -1.8% | 6.09 | 6.06 | -0.5% |
| EC (dS/m) | 0.4 | 0.7 | **73.8%** | 0.7 | 0.8 | 15.4% |
| CEC (meq/100g) | 67.3 | 56.1 | -16.7% | 66.1 | 75.7 | 14.5% |
| OM (%) | 32.2 | 31.8 | -1.1% | 34.2 | 44.3 | **29.6%** |
| Total C (%) | 15.3 | 16.2 | 6.3% | 14.3 | 18.1 | **26.5%** |
| Total N (%) | 0.420 | 0.444 | 5.7% | 0.460 | 0.490 | 5.4% |
| $NO_3$-N (ppm) | 9.0 | 10.9 | **21.4%** | 2.74 | 3.17 | 15.7% |
| $NH_4$-N (ppm) | 8.1 | 7.9 | -2.6% | 22.3 | 26.0 | 16.5% |
| C:N | 36.4 | 36.6 | 0.5% | 31.0 | 37.1 | **20.0%** |
| P (ppm) | 34.6 | 34.0 | -1.5% | 33.2 | 20.9 | **-37.2%** |
| K-av (ppm) | 655 | 570 | -12.9% | 391 | 342 | -12.7% |
| $SO_4$-S (ppm) | 20.3 | 71.2 | **250.1%** | 589 | 593 | 0.7% |
| $CaCO_3$ (%) | 7.20 | 2.98 | **-58.6%** | 0.99 | 1.50 | **52.1%** |
| Zn (ppm) | 0.17 | 0.22 | **25.8%** | 5.96 | 5.70 | -4.4% |
| Fe (ppm) | 27.90 | 13.99 | **-49.9%** | 149 | 142 | -4.4% |
| Mn (ppm) | 1.96 | 1.33 | **-32.0%** | 3723 | 3559 | -4.4% |
| Cn (ppm) | 0.26 | 0.33 | 27.2% | 93086 | 88984 | -4.4% |

### ***CNT affected more taxa of tomato-associated soil microbiomes***

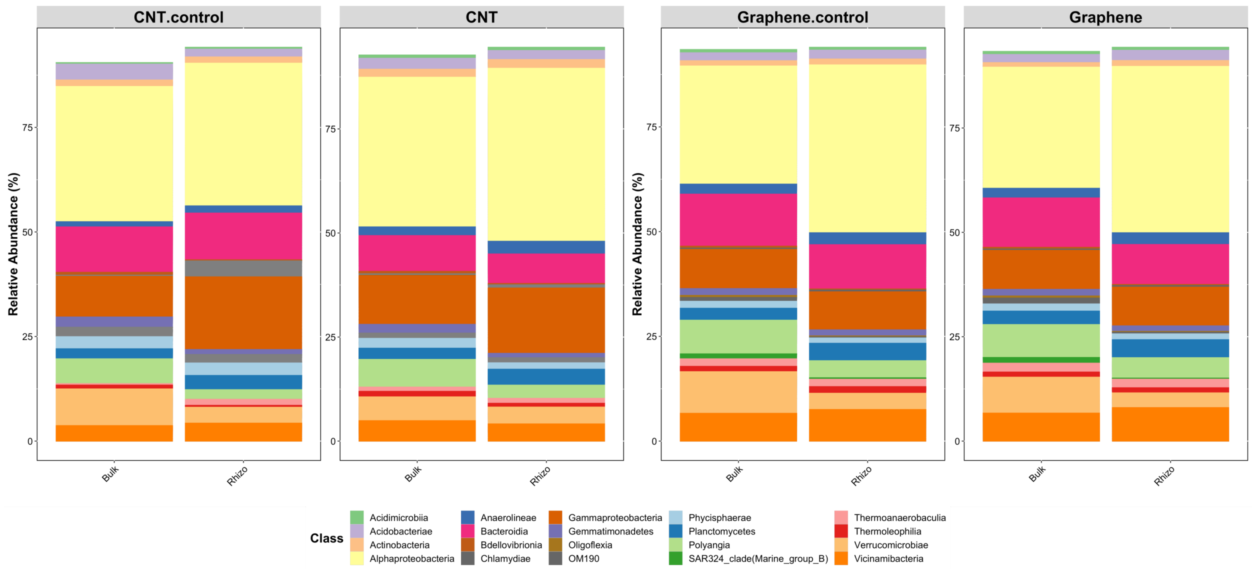


**Figure S2.** Effects of CNT (left) and graphene (right) on the relative abundance of the top 20 classes in the bulk soil and the tomato rhizosphere. Data were averaged among the biological replicates in each experiment.

### *CNT enhanced microbial interactions in the tomato rhizosphere*

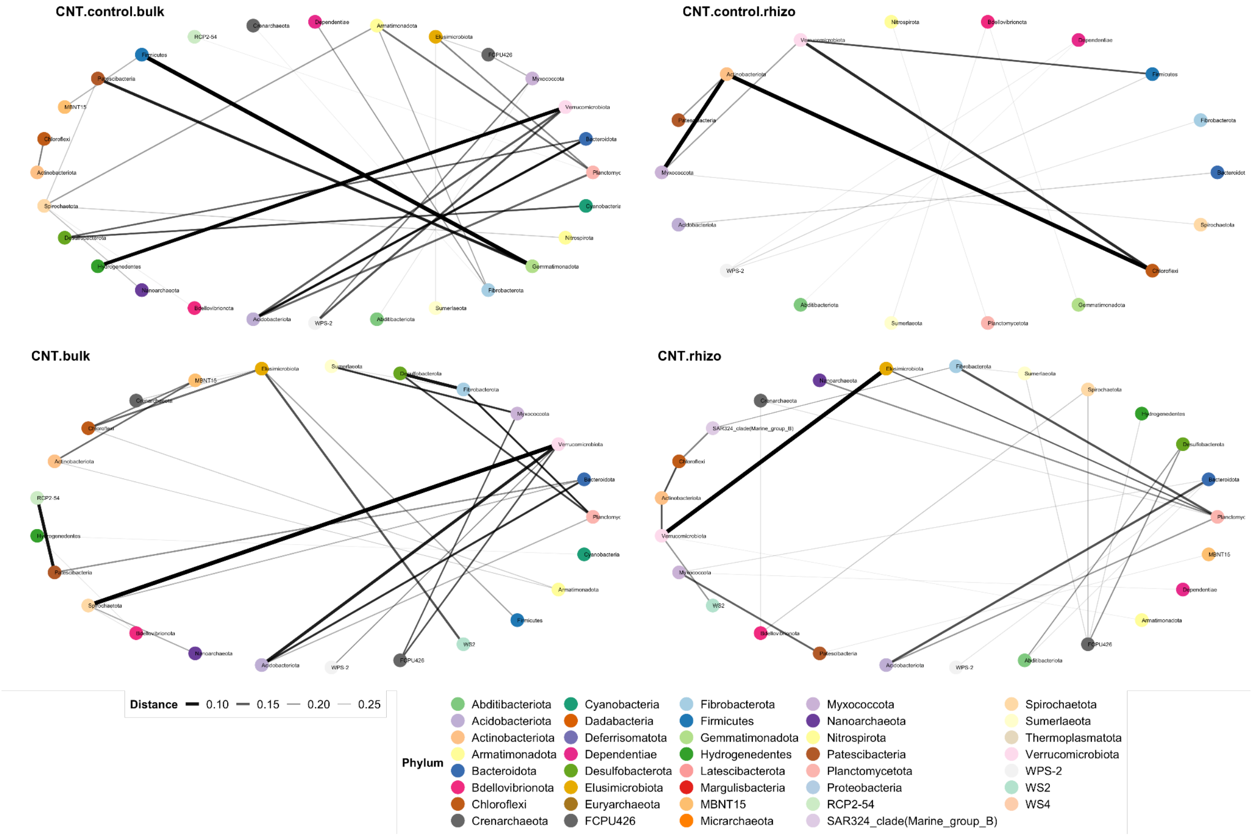


**Figure S3.** Effects of CNT on the class-level microbial network in the bulk soil and the tomato rhizosphere. Networks were calculated based on Bray-Curtis distance with a maximum distance of 0.3.

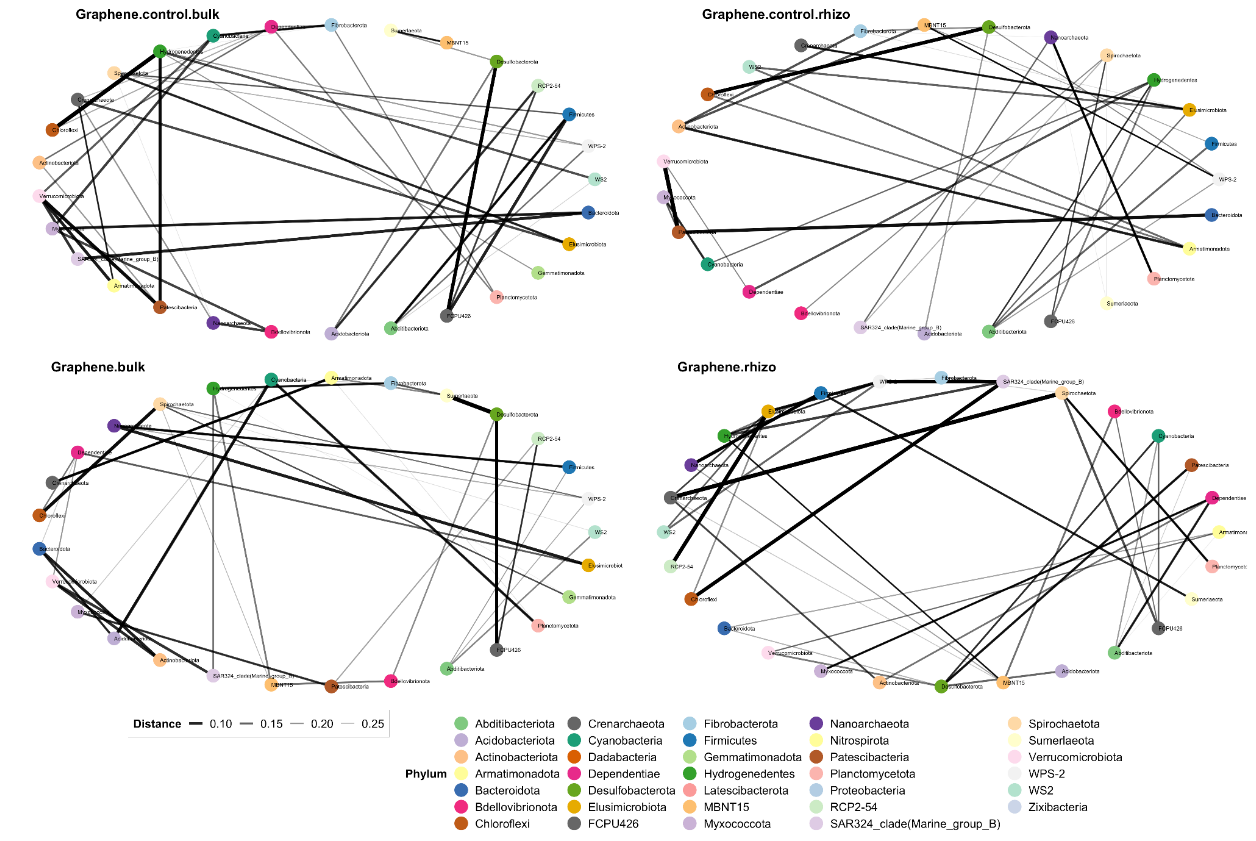


**Figure S4.** Effects of graphene on the class-level microbial network in the bulk soil and the tomato rhizosphere. Networks were calculated based on Bray-Curtis distance with a maximum distance of 0.3.

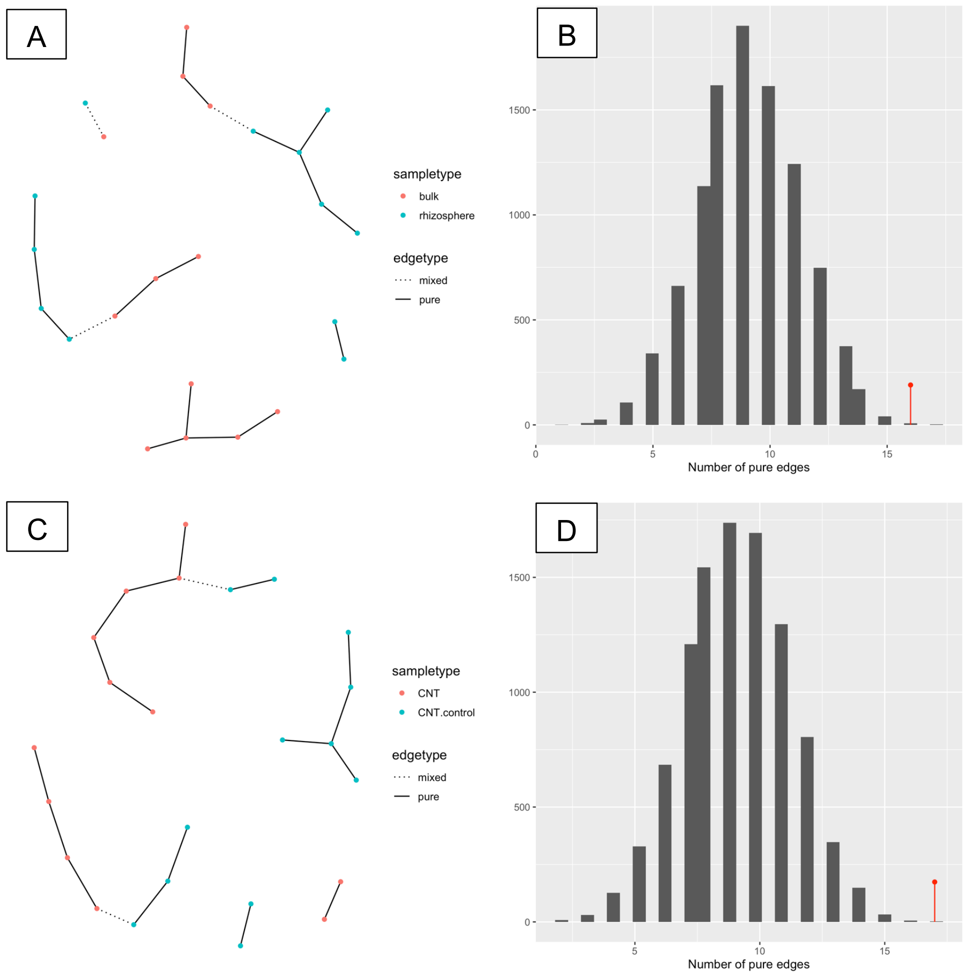


**Figure S5.** In the CNT experiment, soil zone (A, B) and treatment (C, D) are both significant factors shaping microbial network in the bulk soil and the tomato rhizosphere. (A, C) Nearest neighbor (NN) tree constructed on Bray-Curtis distance of agglomerated ASV abundance (to the phylum level) among treatment conditions (nodes). If from the same condition, nodes are connected by solid edges (pure), otherwise they are connected by dashed lines (mixed). Color denotes experimental condition. (B, D) Graph-based permutation test (n = 9999) on the nearest neighbor tree, $p < 0.002$ for both soil zone and treatment.

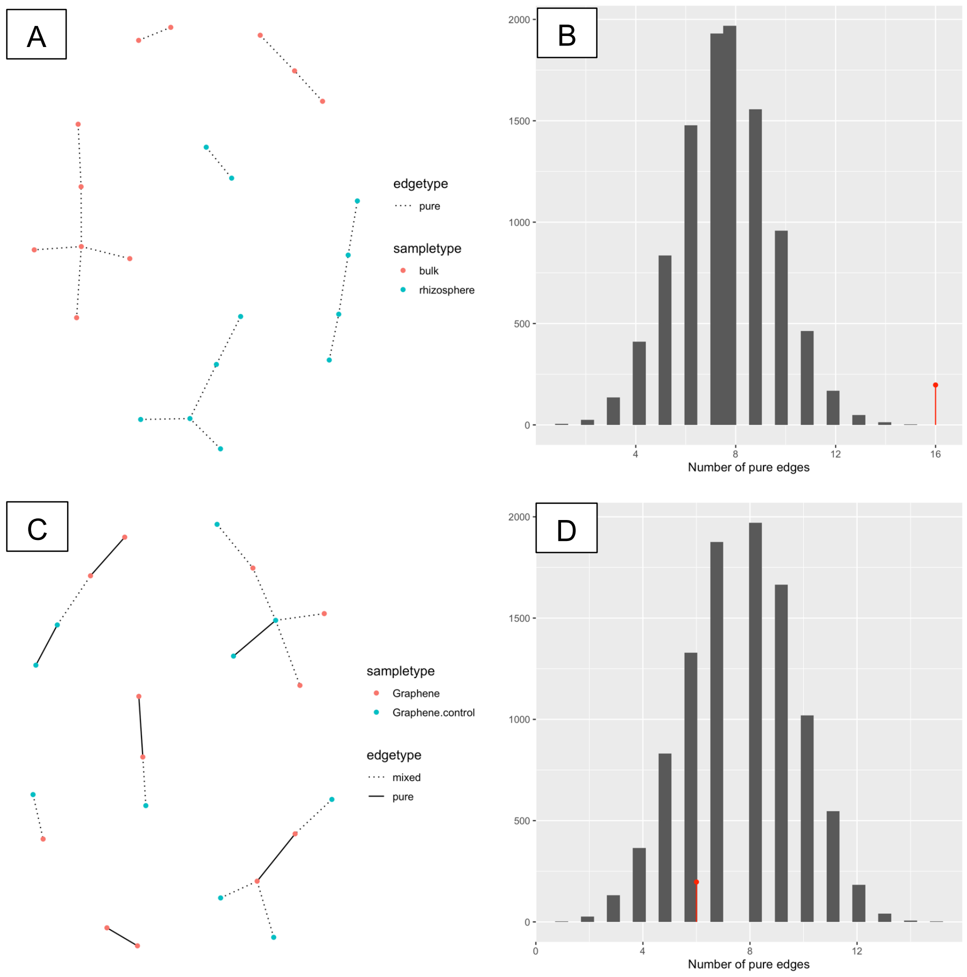


**Figure S6.** In the graphene experiment, soil zone (A, B) but not treatment (C, D) is a significant factor shaping microbial network in the bulk soil and the tomato rhizosphere. (A, C) Nearest neighbor (NN) tree constructed on Bray-Curtis distance of agglomerated ASV abundance (to the phylum level) among treatment conditions (nodes). If from the same condition, nodes are connected by solid edges (pure), otherwise they are connected by dashed lines (mixed). Color denotes experimental condition. (B, D) Graph-based permutation test (n = 9999) on the nearest neighbor tree, $p < 0.001$ for soil zone and $p = 0.864$ for treatment.

***CNT-induced microbial functional changes***

We used PICRUSt2 to infer microbial functions. The resulting NSTI values ranged from 0.135 to 0.258 for all the samples, suggesting a mid-range prediction accuracy typical for soil samples.[3] Rhizosphere samples showed smaller NSTI values than bulk soil samples ($p < 0.002$ in PERMANOVA after controlling treatment conditions), consistent with generally greater microbial diversity in bulk soil. Enriched pathways and modules were identified by DESeq2 in MicrobiomeAnalyst.[4,5] We chose DESeq2 because it uses shrinkage estimation for dispersions and fold changes for improved stability and interpretability of estimates, which enables a more quantitative analysis focused on the strength rather than the mere presence of differential expression.[4]

EcoPlate assay was used to compare community-level metabolism and substrate use patterns in bulk soils at the conclusion of the exposure experiment. After soil sample inoculation, we continuously monitored average well-color development (AWCD) for ~170 hours (Figure S8). AWCD values of the samples reached plateau after 7 days. Therefore, the final AWCD readings were used for calculation and cross-sample comparison. Soil basal respiration was also estimated for bulk soil samples (Figure S9). CNT treatment significantly increased basal respiration in bulk soils ($p < 0.03$ in ANOVA followed by Tukey's HSD). Graphene did not have significant effect on basal soil respiration in bulk soils.

**Table S2.** CNT-induced changes in microbial functional pathways in bulk soil and the tomato rhizosphere. Functional inference was conducted using PICRUSt2.

| Soil zone | Pathway | Total | Expected | Hits | *P*-value | FDR | KO Hits |
|---|---|---|---|---|---|---|---|
| Bulk soil | Nitrogen metabolism | 49 | 2.060 | 10 | 0.0000248 | 0.00392 | K03385<br>K02588<br>K15876<br>K05601<br>K02586<br>K02591<br>K15371<br>K00376<br>K01915<br>K01674 |
| | Amino sugar and nucleotide sugar metabolism | 126 | 5.300 | 13 | 0.00205 | 0.162 | K02795<br>K02796<br>K02794<br>K00844<br>K02564<br>K12454<br>K00849<br>K01209<br>K01809<br>K00963<br>K10012<br>K16011<br>K02377 |
| | Fructose and mannose metabolism | 88 | 3.700 | 10 | 0.00336 | 0.177 | K02795<br>K02796<br>K02794<br>K00844<br>K07046<br>K02770<br>K01809<br>K00895<br>K16011<br>K02377 |
| | Chloroalkane and chloroalkene degradation | 28 | 1.180 | 5 | 0.00548 | 0.216 | K02588<br>K02586 |

| | | | | | | |
|---|---|---|---|---|---|---|
| | | | | | | K02591<br>K00121<br>K00114 |
| Pyruvate metabolism | 86 | 3.620 | 9 | 0.00917 | 0.290 | K00625<br>K00626<br>K01067<br>K00656<br>K01571<br>K01960<br>K01596<br>K01573<br>K01069 |
| Ascorbate and aldarate metabolism | 38 | 1.600 | 5 | 0.020 | 0.484 | K02822<br>K13875<br>K00469<br>K13876<br>K03077 |
| Fatty acid degradation | 39 | 1.640 | 5 | 0.0222 | 0.484 | K00626<br>K06445<br>K00121<br>K01692<br>K00249 |
| Carbon fixation pathways in prokaryotes | 101 | 4.250 | 9 | 0.0245 | 0.484 | K00625<br>K00626<br>K00176<br>K00177<br>K00198<br>K15022<br>K01960<br>K00196<br>K00242 |
| Sulfur metabolism | 74 | 3.110 | 7 | 0.0341 | 0.516 | K01011<br>K00956<br>K08354<br>K08352<br>K00380<br>K00955<br>K01082 |
| Fatty acid metabolism | 60 | 2.520 | 6 | 0.0384 | 0.516 | K00626<br>K06445 |

| | | | | | | | |
|---|---|---|---|---|---|---|---|
| | | | | | | | K16363<br>K01692<br>K00249<br>K01716 |
| | Porphyrin and chlorophyll metabolism | 76 | 3.200 | 7 | 0.0387 | 0.516 | K04040<br>K03428<br>K04037<br>K04038<br>K04039<br>K10960<br>K03403 |
| | Methane metabolism | 147 | 6.180 | 11 | 0.0427 | 0.516 | K00625<br>K12234<br>K03388<br>K11212<br>K00121<br>K00198<br>K15022<br>K00196<br>K03390<br>K11261<br>K02203 |
| | Biosynthesis of amino acids | 223 | 9.380 | 15 | 0.0446 | 0.516 | K01914<br>K11358<br>K05825<br>K05822<br>K05823<br>K13853<br>K17462<br>K14155<br>K01960<br>K01243<br>K01915<br>K02502<br>K03785<br>K14682<br>K02203 |
| Rhizosphere | Toluene degradation | 38 | 1.63 | 15 | 9.70E-12 | 1.53E-09 | K16242<br>K16243<br>K16244 |

| | | | | | | |
|---|---|---|---|---|---|---|
| | | | | | | K16245<br>K16246<br>K16249<br>K00141<br>K05797<br>K07540<br>K07543<br>K07545<br>K07547<br>K07548<br>K07549<br>K07550 |
| Degradation of aromatic compounds | 171 | 7.35 | 26 | 5.95E-09 | 4.70E-07 | K04072<br>K16242<br>K16243<br>K16244<br>K16245<br>K16246<br>K07537<br>K16249<br>K00141<br>K10216<br>K05712<br>K05783<br>K04108<br>K07540<br>K18074<br>K18076<br>K07536<br>K16050<br>K07543<br>K07545<br>K07547<br>K07548<br>K07549<br>K07550<br>K16049<br>K05550 |
| Benzoate degradation | 82 | 3.53 | 17 | 3.22E-08 | 1.69E-06 | K04098<br>K16242 |

| | | | | | | |
|---|---|---|---|---|---|---|
| | | | | | | K16243<br>K16244<br>K16245<br>K16246<br>K07537<br>K16249<br>K04110<br>K10216<br>K05783<br>K10221<br>K04108<br>K01615<br>K07536<br>K05550<br>K04100 |
| Steroid degradation | 9 | 0.387 | 4 | 0.000348 | 0.0115 | K16050<br>K05898<br>K16049<br>K15982 |
| Chlorocyclohexane and chlorobenzene degradation | 33 | 1.42 | 7 | 0.000389 | 0.0115 | K04098<br>K16242<br>K16243<br>K16244<br>K16245<br>K16246<br>K16249 |
| Methane metabolism | 147 | 6.32 | 16 | 0.000438 | 0.0115 | K00196<br>K01007<br>K03389<br>K05979<br>K03390<br>K11781<br>K11780<br>K16792<br>K00148<br>K11212<br>K12234<br>K00198<br>K15022<br>K03388 |

| | | | | | | |
|---|---|---|---|---|---|---|
| Phenylalanine metabolism | 76 | 3.27 | 10 | 0.0013 | 0.0294 | K18277<br>K03841<br>K01912<br>K10775<br>K11358<br>K05712<br>K02614<br>K02610<br>K02611<br>K02612<br>K02609<br>K02613<br>K00141 |
| Aminobenzoate degradation | 35 | 1.51 | 5 | 0.0157 | 0.309 | K04110<br>K10221<br>K04108<br>K04100<br>K05928 |
| Ubiquinone and other terpenoid-quinone biosynthesis | 50 | 2.15 | 6 | 0.0189 | 0.332 | K03182<br>K09833<br>K09834<br>K18534<br>K12073 |
| Carotenoid biosynthesis | 28 | 1.2 | 4 | 0.030 | 0.474 | K09835<br>K02293<br>K14605<br>K14606 |
| Xylene degradation | 32 | 1.38 | 4 | 0.0462 | 0.659 | K00141<br>K10216<br>K05783<br>K05550 |

**Table S3.** CNT-induced changes in microbial functional modules in bulk soil and the tomato rhizosphere. Functional inference was conducted using PICRUSt2.

| Soil zone | Module | Total | Expected | Hits | *P*-value | FDR | KO Hits (relative change) |
|---|---|---|---|---|---|---|---|
| Bulk soil | Nitrogen fixation, nitrogen => ammonia | 4 | 0.170 | 3 | 0.000282 | 0.067 | K02588<br>K02586<br>K02591 |
| | Assimilatory sulfate reduction, sulfate => $H_2S$ | 10 | 0.424 | 3 | 0.00704 | 0.830 | K00956<br>K00380<br>K00955 |
| | Methanogenesis, acetate => methane | 13 | 0.552 | 3 | 0.0153 | 0.944 | K00625<br>K03388<br>K03390 |
| | $F_{420}$ biosynthesis | 6 | 0.255 | 2 | 0.0238 | 0.944 | K12234<br>K11212 |
| | Incomplete reductive citrate cycle, acetyl-CoA => oxoglutarate | 17 | 0.721 | 3 | 0.0323 | 0.944 | K00176<br>K00177<br>K01960 |
| | Nucleotide sugar biosynthesis, glucose => UDP-glucose | 7 | 0.297 | 2 | 0.0324 | 0.944 | K00844<br>K00963 |
| | Cysteine biosynthesis, methionine => cysteine | 7 | 0.297 | 2 | 0.0324 | 0.944 | K17462<br>K01243 |
| | Ascorbate degradation, ascorbate => D-xylulose-5P | 7 | 0.297 | 2 | 0.0324 | 0.944 | K02822<br>K03077 |
| | Methanogenesis, CO2 => methane | 19 | 0.806 | 3 | 0.0433 | 0.944 | K03388<br>K03390<br>K11261 |
| | beta-Oxidation | 19 | 0.806 | 3 | 0.0433 | 0.944 | K06445<br>K01692<br>K00249 |
| Rhizosphere | Benzene degradation, benzene => catechol | 6 | 0.303 | 6 | 1.34E-08 | 2.42E-06 | K16242<br>K16243<br>K16244<br>K16245<br>K16246<br>K16249 |
| | Toluene degradation, anaerobic, toluene => benzyl-CoA | 9 | 0.454 | 7 | 2.05E-08 | 2.42E-06 | K07540<br>K07543<br>K07545 |

| | | | | | | |
|---|---|---|---|---|---|---|
| | | | | | | K07547<br>K07548<br>K07549<br>K07550 |
| Tocopherol biosynthesis | 5 | 0.252 | 4 | 2.87E-05 | 0.00226 | K05928<br>K09833<br>K09834<br>K18534 |
| $F_{420}$ biosynthesis | 6 | 0.303 | 4 | 8.28E-05 | 0.00489 | K11781<br>K11780<br>K11212<br>K12234 |
| Methanogenesis, methanol => methane | 9 | 0.454 | 3 | 0.00831 | 0.392 | K03389<br>K03390<br>K03388 |
| Benzoate degradation, benzoate => catechol / methylbenzoate => methylcatechol | 4 | 0.202 | 2 | 0.0141 | 0.476 | K05783<br>K05550 |
| Terephthalate degradation, terephthalate => 3,4-dihydroxybenzoate | 4 | 0.202 | 2 | 0.0141 | 0.476 | K18074<br>K18076 |
| Methanogenesis, acetate => methane | 13 | 0.656 | 3 | 0.0245 | 0.641 | K03389<br>K03390<br>K03388 |
| Methanogenesis, methylamine/dimethylamine/trimethylamine => methane | 13 | 0.656 | 3 | 0.0245 | 0.641 | K03389<br>K03390<br>K03388 |
| beta-Carotene biosynthesis, GGAP => beta-carotene | 6 | 0.303 | 2 | 0.033 | 0.779 | K09835<br>K02293 |
| Reductive pentose phosphate cycle, ribulose-5P => glyceraldehyde-3P | 7 | 0.353 | 2 | 0.0447 | 0.96 | K00150<br>K01602 |

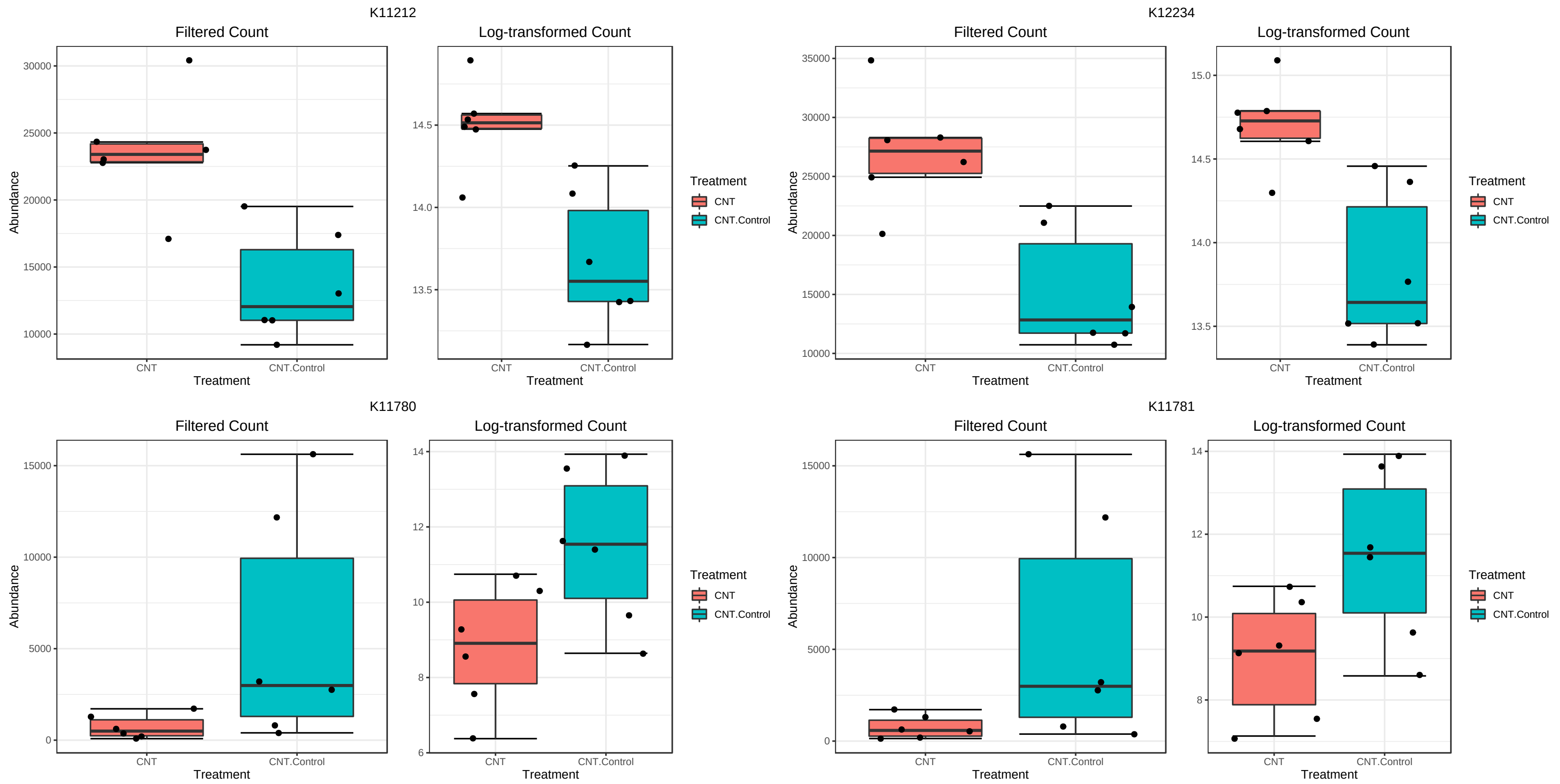


**Figure S7.** CNT effects on $F_{420}$ biosynthesis in the tomato rhizosphere. K11212: *cofD* gene encoding 2-phospho-L-lactate transferase; K12234: *cofE* gene encoding coenzyme F420:L-glutamate ligase; K11780: *cofG* encoding 7,8-didemethyl-8-hydroxy-5-deazariboflavin synthase; K11781: *cofH* gene encoding 5-amino-6-(D-ribitylamino)uracil-L-tyrosine 4-hydroxyphenyl transferase.[6]

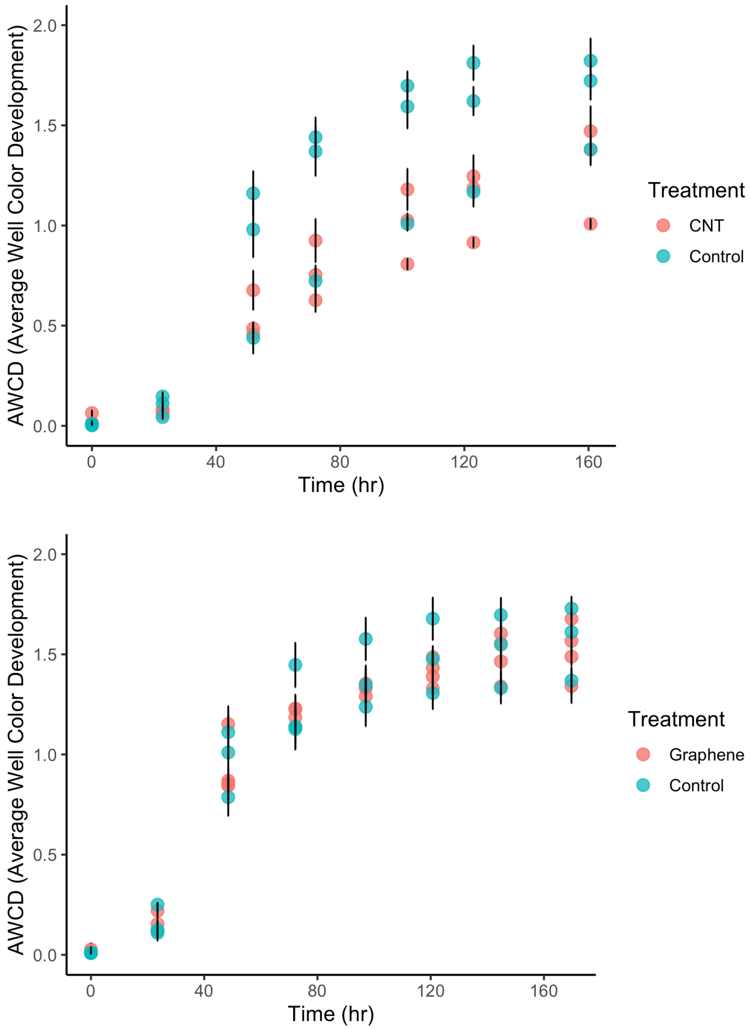


**Figure S8.** AWCD of bulk soils harvested by the end of the CNT or graphene experiment. Data points represent average across 3 technical replicates and 3-4 biological replicates; bars represent standard deviation.

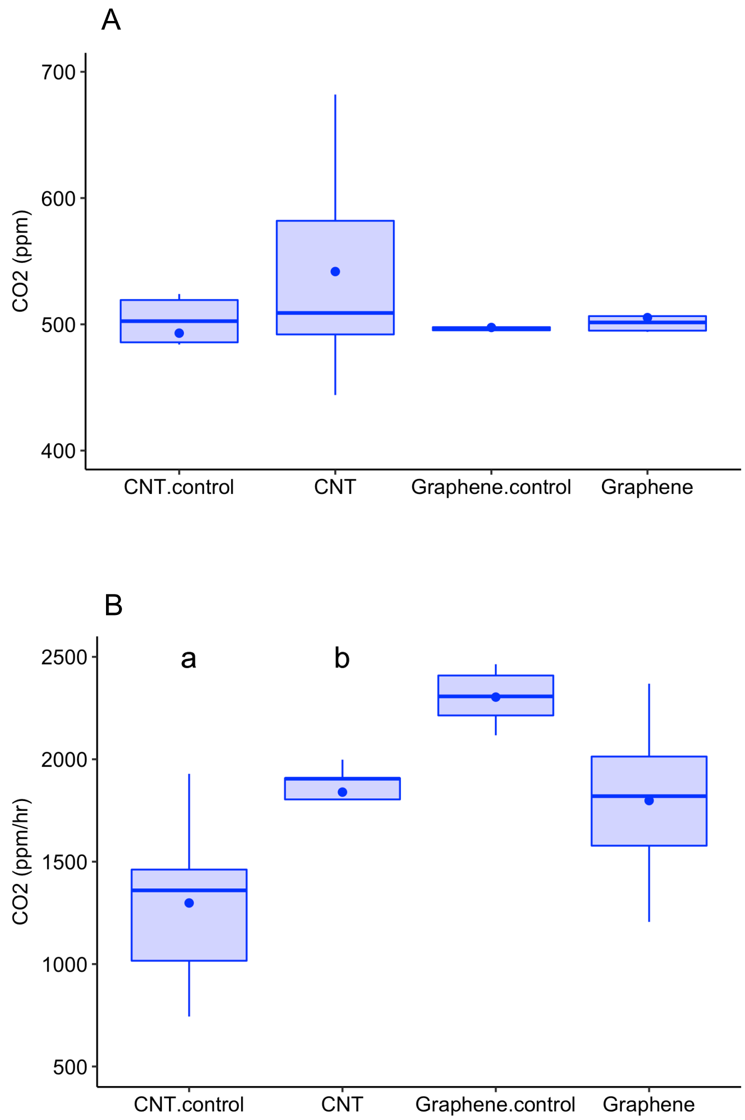


**Figure S9.** (A) Ambient $CO_2$ measured at the time of soil respiration assessment. No significant difference was observed among measurements. (B) Soil basal respiration rate (hourly $CO_2$ generation) in the treated and untreated soils. Letters indicate statistical significance ($p < 0.03$ in ANOVA followed by Tukey's HSD).

### References


1. S.R. Olsen, Estimation of available phosphorus in soils by extraction with sodium bicarbonate (No. 939), US Department of Agriculture, 1954.

2. L.E. Allison and C.D. Moodie, Carbonate. Methods of Soil Analysis: Part 2 Chemical and Microbiological Properties, 1965, 9, pp.1379-1396.

3. G.M. Douglas, V.J. Maffei, J.R. Zaneveld, S.N. Yurgel, J.R. Brown, C.M. Taylor, C. Huttenhower and M.G. Langille, PICRUSt2 for prediction of metagenome functions, *Nat. Biotechnol.*, 2020, **38**, 685-688.

4. M.I. Love, W. Huber and S. Anders, Moderated estimation of fold change and dispersion for RNA-seq data with DESeq2, *Genome Biol.*, 2014, **15**, 1-21.

5. J. Chong, P. Liu, G. Zhou and J. Xia, Using MicrobiomeAnalyst for comprehensive statistical, functional, and meta-analysis of microbiome data, *Nat. Protoc.*, 2020, **15**, 799-821.

6. R. Grinter and C. Greening, Cofactor F420: An expanded view of its distribution, biosynthesis and roles in bacteria and archaea, *FEMS Microbiol. Rev.*, 2021, **45**, fuab021.
